# Identification of novel protein lysine acetyltransferases in *Escherichia coli*

**DOI:** 10.1101/408930

**Authors:** David G. Christensen, Jesse G. Meyer, Jackson T. Baumgartner, Alexandria K. D’Souza, William C. Nelson, Samuel H. Payne, Misty L. Kuhn, Birgit Schilling, Alan J. Wolfe

## Abstract

Post-translational modifications, such as Nε-lysine acetylation, regulate protein function. Nε-lysine acetylation can occur either non-enzymatically or enzymatically. The non-enzymatic mechanism uses acetyl phosphate (AcP) or acetyl coenzyme A (AcCoA) as acetyl donors to modify an Nε-lysine residue of a protein. The enzymatic mechanism uses Nε-lysine acetyltransferases (KATs) to specifically transfer an acetyl group from AcCoA to Nε-lysine residues on proteins. To date, only one KAT (YfiQ, also known as Pka and PatZ) has been identified in *E. coli*. Here, we demonstrate the existence of 4 additional *E. coli* KATs: RimI, YiaC, YjaB, and PhnO. In a genetic background devoid of all known acetylation mechanisms (most notably AcP and YfiQ) and one deacetylase (CobB), overexpression of these putative KATs elicited unique patterns of protein acetylation. We mutated key active site residues and found that most of them eliminated enzymatic acetylation activity. We used mass spectrometry to identify and quantify the specificity of YfiQ and the four novel KATs. Surprisingly, our analysis revealed a high degree of substrate specificity. The overlap between KAT-dependent and AcP-dependent acetylation was extremely limited, supporting the hypothesis that these two acetylation mechanisms play distinct roles in the post-translational modification of bacterial proteins. We further showed that these novel KATs are conserved across broad swaths of bacterial phylogeny. Finally, we determined that one of the novel KATs (YiaC) and the known KAT (YfiQ) can negatively regulate bacterial migration. Together, these results emphasize distinct and specific non-enzymatic and enzymatic protein acetylation mechanisms present in bacteria.

**Importance:** Nε-lysine acetylation is one of the most abundant and important post-translational modifications across all domains of life. One of the best-studied effects of acetylation occurs in eukaryotes, where acetylation of histone tails activates gene transcription. Although bacteria do not have true histones, Nε-lysine acetylation is prevalent; however, the role of these modifications is mostly unknown. We constructed an *E. coli* strain that lacked both known acetylation mechanisms to identify four new Nε-lysine acetyltransferases (RimI, YiaC, YjaB, and PhnO). We used mass spectrometry to determine the substrate specificity of these acetyltransferases. Structural analysis of selected substrate proteins revealed site-specific preferences for enzymatic acetylation that had little overlap with the preferences of the previously reported acetyl-phosphate non-enzymatic acetylation mechanism. Finally, YiaC and YfiQ appear to regulate flagellar-based motility, a phenotype critical for pathogenesis of many organisms. These acetyltransferases are highly conserved and reveal deeper and more complex roles for bacterial post-translational modification.

## Introduction

During Nε-lysine acetylation, an acetyl group is added to the epsilon amino group of a lysine residue of a protein, which neutralizes the positive charge and increases the size of the sidechain. In eukaryotes, the effects of protein acetylation have been well described, historically and most fully in the context of histone tail acetylation, which regulates eukaryotic transcription. Over the past decade, it has become clear that many non-histone proteins are also Nε-lysine acetylated and that this post-translational modification is similarly abundant in archaea and bacteria (1-4).

In *Escherichia coli*, two mechanisms of Nε-lysine acetylation are known. The predominant mechanism is a non-enzymatic donation of an acetyl group from the high-energy central metabolite acetyl-phosphate (AcP) onto a susceptible lysine of a protein (5, 6). AcP is the intermediate of the phosphotransferase (Pta) – acetate kinase (AckA) pathway that interconverts acetyl-CoA (AcCoA), inorganic phosphate, and ADP to acetate, CoA, and ATP. Alternatively, an Nε-lysine acetyltransferase (KAT) can catalyze acetylation of a specific lysine using AcCoA as the acetyl donor. In *E. coli*, only one KAT has been discovered to date, YfiQ (also called Pka and PatZ) (7-9). This KAT was first discovered in *Salmonella enterica*, where it is called Pat; elegant studies showed that it inactivates acetyl-CoA synthetase (Acs) by acetylation, preventing acetate assimilation (8). It is conserved in many other bacterial species including *E. coli* (4). Global acetylation profiles of Δ*yfiQ* mutants provide evidence that YfiQ might acetylate targets in addition to Acs (5, 6). Other studies report that YfiQ can acetylate RNA polymerase, RNase R, RNase II, and DnaA on lysines that alter function (10-13). Transcription of *yfiQ* depends on activation by cyclic AMP (cAMP)-bound catabolite activator protein (CAP; also known as the cAMP receptor protein or CRP) (14). During growth in minimal glucose medium, it is upregulated in stationary phase. However, another report using *E. coli* strain MG1655 in YT medium showed that YfiQ protein levels are reduced in stationary phase (15). Thus, there is more to learn about the regulation of YfiQ and consequence of acetylation.

YfiQ is just one member of the Gcn5-related *N*-acetyltransferase (GNAT) family. GNATs acetylate a broad range of substrates, including antibiotics, polyamines, amino acids, nucleotides, tRNAs, proteins, and peptides (4, 16). The *E. coli* K-12 genome encodes 26 genes whose products are annotated as GNATs, of which only about half have annotated functions (4, 17, 18). Therefore, we investigated whether other *E. coli* GNATs beyond YfiQ could function as KATs.

Using an *E. coli* strain that lacked both known acetylation mechanisms (YfiQ and AcP), we found four GNATs that increased relative site-specific acetylation levels on proteins *in vivo* as one would expect for a KAT. By western immunoblot and mass spectrometric analyses, we demonstrated that these four GNATs facilitate increased Nε-lysine acetylation and identified their cognate substrates. This increased acetylation was lost upon mutation of conserved catalytic or active site residues found in other known GNATs. We conclude that *E. coli* encodes multiple KATs that exhibit substrate specificities that differ from non-enzymatic acetylation by AcP.

## Results

### Identification of putative uncharacterized KATs

If acetylation in *E. coli* depends solely on the two known mechanisms of protein acetylation, then we would be unable to detect acetylation in a mutant that lacks: 1) the only known *E. coli* acetyltransferase YfiQ, and 2) the ability to generate AcP, either by deleting Pta or the entire Pta-AckA pathway. However, anti-acetyllysine western blot analysis of a strain that lacked both of these mechanisms (Δ*ackA pta yfiQ*) revealed that residual acetylation remained (**Fig. S1**). We therefore sought the mechanism(s) behind this residual AcP-and YfiQ-independent acetylation and hypothesized that this residual acetylation activity could be attributed to uncharacterized KATs. Since YfiQ is a member of the GNAT family of proteins, and *E. coli* contains 25 other members of this family, we tested whether these proteins had KAT activity.

To determine whether these GNATs have KAT activity, we compared acetylation profiles of strains overexpressing each of the GNATs via anti-acetyllysine western blot. We used a Δ*pta yfiQ acs cobB* background to enhance the signal-to-noise ratio, which we refer to as the acetylation “gutted” strain. This strain reduces background acetylation levels from AcP and YfiQ (Δ*pta yfiQ*), while ensuring that residual acetylation that occurs would not be reversed by the CobB deacetylase (Δ*cobB*). Acs was also deleted, as it has been reported to acetylate the chemotaxis response regulator CheY (19). Furthermore, YfiQ regulates Acs activity and loss of that control can have a detrimental effect on growth (20). As with the Δ*pta ackA yfiQ* mutant (**Fig. S1**), the gutted strain (Δ*pta yfiQ acs cobB*) exhibited only limited acetylation (**Fig. 1**). To validate that this strain behaved as expected and hyper-acetylated specific lysine sites with the known KAT, YfiQ, we first compared YfiQ overexpression in a gutted strain that expresses the YfiQ substrate Acs (Δ*pta yfiQ cobB*, Acs^+^) to a gutted strain that does not express Acs (Δ*pta yfiQ cobB*, Acs^-^). Indeed, by anti-acetyllysine western blot, we observed an acetylated band in the gutted Acs^+^ strain that was absent in the gutted Acs^-^ strain (**Fig. 2** and **S2**, in **Fig. 2,** compare lane 1 [positive control] and lane 6 [YfiQ]).

**FIGURE 1.**
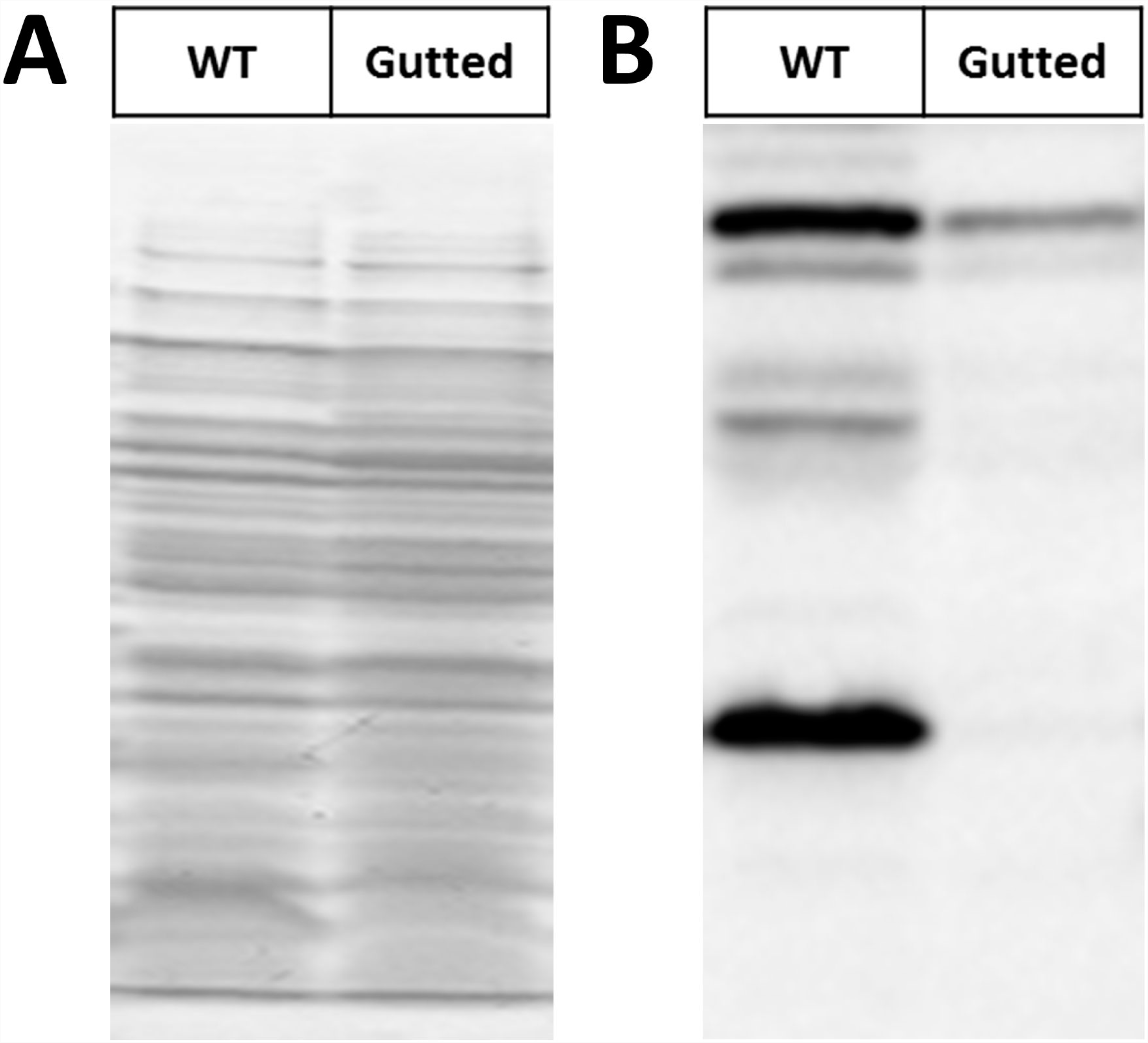
Inactivation of the two known acetylation mechanisms in *E. coli* eliminates the majority of acetylation. Wild-type (WT) *E. coli* (strain BW25113) and an isogenic Δ*pta yfiQ acs cobB* mutant (Gutted) were aerated in TB7 supplemented with 0.4% glucose for 10 hours. Whole cell lysates were analyzed (A) by Coomassie blue-stained SDS-polyacrylamide gel to ensure equivalent loading and (B) by anti-acetyllysine western blot.

**FIGURE 2.**
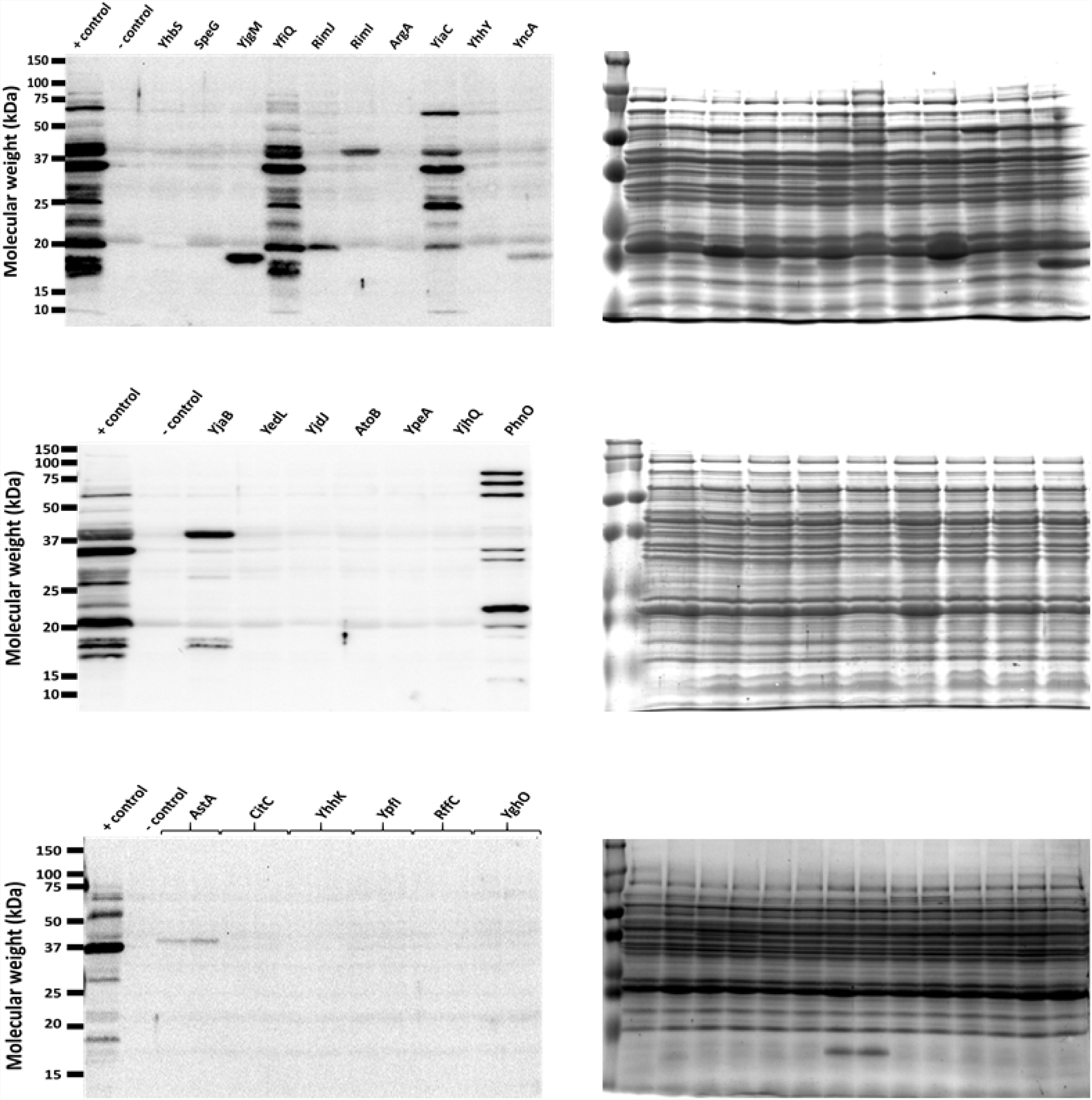
Overexpression of five GNAT family members results in altered lysine acetylation patterns by anti-acetyllysine western blot. The gutted strain (BW25113 Δ*pta yfiQ acs cobB*) was transformed with the pCA24n vector control (negative (-) control) or pCA24n containing the indicated genes under an IPTG inducible promoter (69). As a positive (+) control, an isogenic strain that retained the WT allele of *acs* (Δ*pta yfiQ cobB*) was transformed with pCA24n containing YfiQ. The resulting strains were aerated in TB7 supplemented with 0.4% glucose, 50 μM IPTG, and 25 μg/mL chloramphenicol for 10 hours. Whole cell lysates were analyzed (right panels) by Coomassie blue-stained SDS-polyacrylamide gel to ensure equivalent loading and (left panels) by anti-acetyllysine western blot. Note that the band in RimJ was not reproducible. The positive control contains one additional YfiQ-dependent band around 72 kDa, which corresponds to Acs (82). YncA and AstA each produce an acetylated band that can be observed in the Coomassie stained gel at the expected molecular weight of these proteins.

Upon induction of each of the 25 GNAT family members in the gutted strain, overexpression of four GNATs (Aat, ElaA, YiiD, and YafP) inhibited growth. For the 21 strains that did grow, only 8 of the putative GNATs – plus YfiQ-resulted in the appearance of one or more acetylated protein band(s) (**Fig. 2**). Induction of RimI, YiaC, YjaB, YjgM, and PhnO expression produced reproducible acetylated protein band(s) (**Fig. S3**); in contrast, induction of RimJ did not (data not shown). Induction of YncA (17 kDa) and AstA (38.5 kDa) each produced a single acetylated band that migrated consistent with their expected molecular masses, suggesting acetylation of the proteins themselves. We selected RimI, YiaC, YjaB and PhnO for further assessment of their ability to function as KATs.

### Mutation of conserved catalytic amino acids inactivates RimI, YiaC, YjaB, and PhnO

To determine whether these GNATs directly acetylated protein targets, we mutated a few residues that could act as general acids/bases in the reaction or could be important for protein substrate recognition. Acetyltransferases acetylate their substrates using a general acid/base chemical mechanism. Typically, a glutamate (E) or water molecule within a KAT active site acts as the general base by deprotonating the amino group of the substrate. This permits the nitrogen of the amino group to attack the carbonyl carbon of the acetyl group of AcCoA, and results in an acetylated product and CoA anion. An amino acid such as tyrosine (Y) then acts as the general acid to reprotonate the thiolate of CoA (21). To select amino acids for mutagenesis, we compared the putative GNAT sequences using the protein structure prediction tool, Phyre2 (22) and then generated the following mutants based on this analysis: YiaC (F70A, Y115A), YjaB (Y117A, Y117F), RimI (Y115A), and PhnO (E78A, Y128A). Plasmids carrying the mutant alleles were introduced into the gutted strain, and the cell lysates were analyzed for successful expression of the mutant proteins and for acetylation. All putative KAT variants were detected at comparable levels by anti-His6 western blot, except YjaB Y117A, whose levels were clearly reduced relative to its wild-type isoform (**Fig. 3A, B**). Overexpression of all tyrosine and glutamate mutants of YiaC, YjaB, RimI, and PhnO eliminated the acetylation signal produced by the wild-type isoforms (**Fig. 3C, D**). However, the YiaC F70A mutant did not completely lose activity as it produced the same acetylated bands as the wild-type isoform, but with reduced intensity.

**FIGURE 3.**
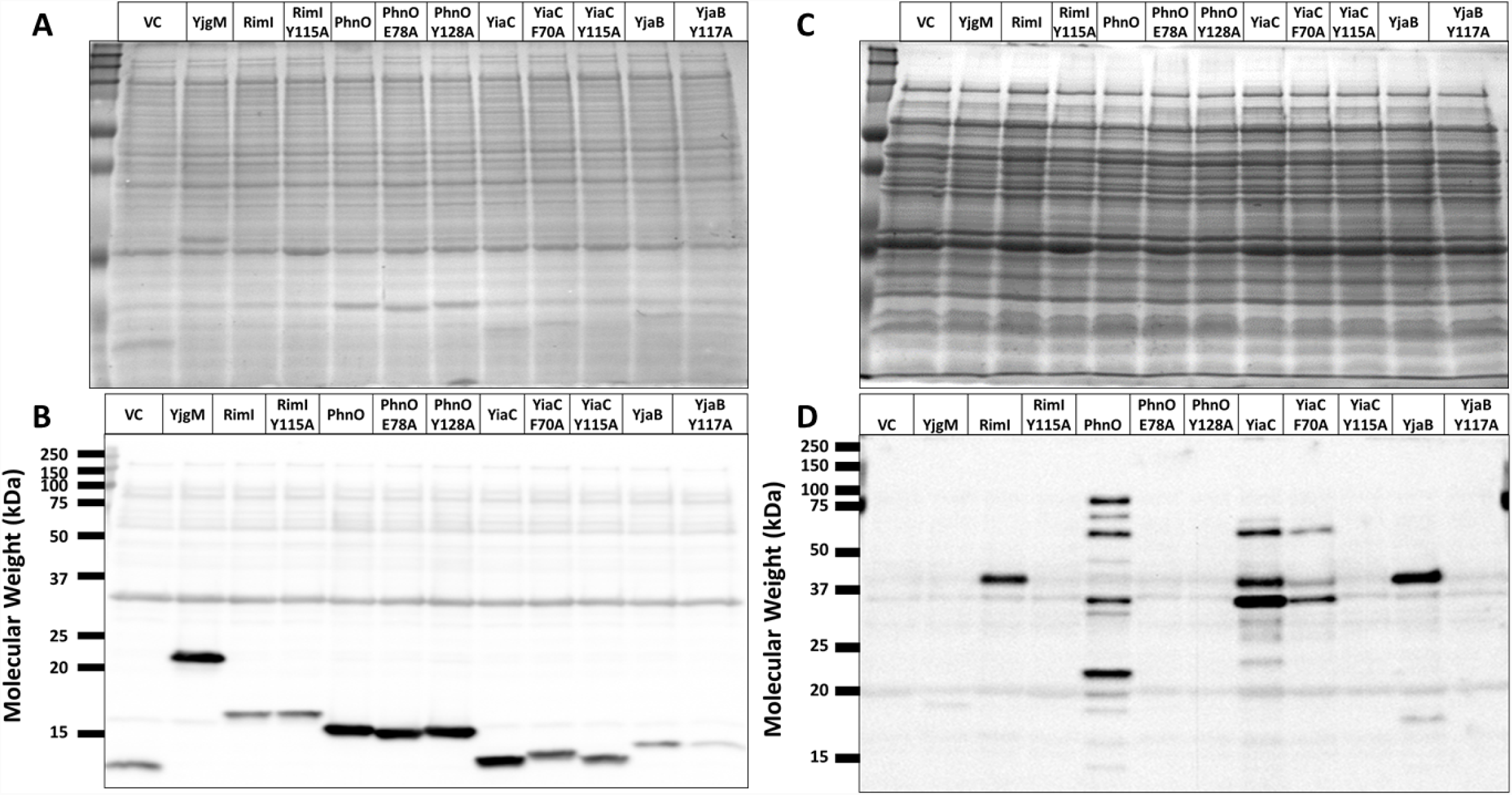
Mutation of conserved catalytic amino acids prevents RimI, PhnO, YjaB, and YiaC-dependent acetylation. The gutted strain (BW25113 Δ*pta yfiQ acs cobB*) was transformed with the pCA24n vector control, pCA24n carrying the wild-type allele for each putative KAT, or mutant alleles for each putative KAT with alanine substitutions of the indicated residues. The resulting strains were grown in TB7 supplemented with 0.4% glucose, 100 μM IPTG, and 25 μg/mL chloramphenicol for 8 hours. Crude lysates harvested after 4 hours were analyzed for expression of the KAT proteins. Whole cell lysates harvested after 8 hours were analyzed for acetylation. Coomassie stained SDS-PAGE gels (A, C) served as loading controls for anti-His (B) and an anti-acetyllysine (D) western blots.

Because the amount of the YjaB Y117A protein was reduced relative to wild-type YjaB in the anti-His6 western blot, we mutated this residue to phenylalanine (Y117F) to determine if soluble expression of this mutant improved. This Y-to-F mutation removes the hydroxyl group involved in re-protonation of CoA but retains the phenyl ring. The YjaB Y117F mutant showed similar soluble expression levels compared to WT and a decreased acetylation signal similar to that of the other tyrosine mutants (**Fig. S4**). Overall, these data provided very strong evidence that RimI, YiaC, YjaB, and PhnO function as KATs.

### Identification of putative KAT substrate proteins by mass spectrometry

Given the evidence that these four GNATs function as KAT enzymes, we sought to identify their substrate proteins and the amino acids that they acetylate. We used acetyllysine enrichment and mass spectrometry for unbiased identification and quantification of acetylation sites as described previously (**Fig. 4A**) (5, 23, 24). Proteome samples were isolated from the *E. coli* strains overexpressing RimI, YiaC, YjaB, and PhnO, as well as the known acetyltransferase YfiQ as a positive control and empty vector as a negative control. Using the standard workflow with trypsin digestion, we identified 1240 unique acetylation sites on 586 unique proteins (**Fig. 4B, Table S1A**). To increase the protein sequence coverage and therefore quantifiable acetylation sites, we performed the same experiments in parallel but substituted trypsin for a complementary protease, GluC, (25-27), which expanded the total number of identifications by nearly 25% to 1539 unique acetylation sites on 668 proteins (**Fig. 4B, Table S1A**).

To determine the set of acetylation sites regulated by these novel KATs and YfiQ, we applied stringent filters to the quantitative comparisons between the overexpression samples and controls (q-value < 0.01 and log_2_ (FC) ≥ 2, which is a ≥ 4-fold increase), resulting in a total of 818 acetylation sites on 434 proteins whose acetylation increased with overexpression of at least one KAT (**Fig. 4B**). These altered acetylation site levels were not driven by proteome remodeling, as only a handful of proteins were altered due to overexpression of any KAT (**Table S1B**). Again, the additional data from GluC digestion proved complementary, revealing 122 additional significantly increased acetylation sites (**Fig. 4C**). The acetylation sites, their fold-increase, and the overlap between these putative KATs and YfiQ is shown as a heat map in **Figure 4D**. As expected, the known acetyltransferase, YfiQ, acetylated the most lysines, with a total of 649 sites with significantly enhanced acetylation on 364 proteins (**Table 1**). YiaC and YjaB overexpression resulted in fewer, yet substantial, numbers of significantly increased acetylation of sites/proteins (391/251 and 171/128, respectively). Overexpression of RimI and PhnO elicited the fewest changes, each acetylating fewer than 20 sites. It should be noted that we observed many more acetylated proteins by mass spectrometry when compared to the number of bands we obtained via western blot analysis (**Fig. 1B**). Mass spectrometry will detect site-specific acetylated peptides with greater sensitivity than western blot as previously shown (5, 23). Additionally, different acetylated proteins may migrate together on a gel and result in the appearance of only one band on a western blot.

**Table 1.** Number of proteins and lysine residues with significantly increased acetylation upon overexpression of KATs

| KAT | # of Unique proteins acetylated | # of Unique Lysines acetylated |
| --- | --- | --- |
| YfiQ | 364 | 649 |
| YjaB | 128 | 171 |
| YiaC | 251 | 391 |
| RimI | 11 | 14 |
| PhnO | 10 | 11 |

**FIGURE 4.**
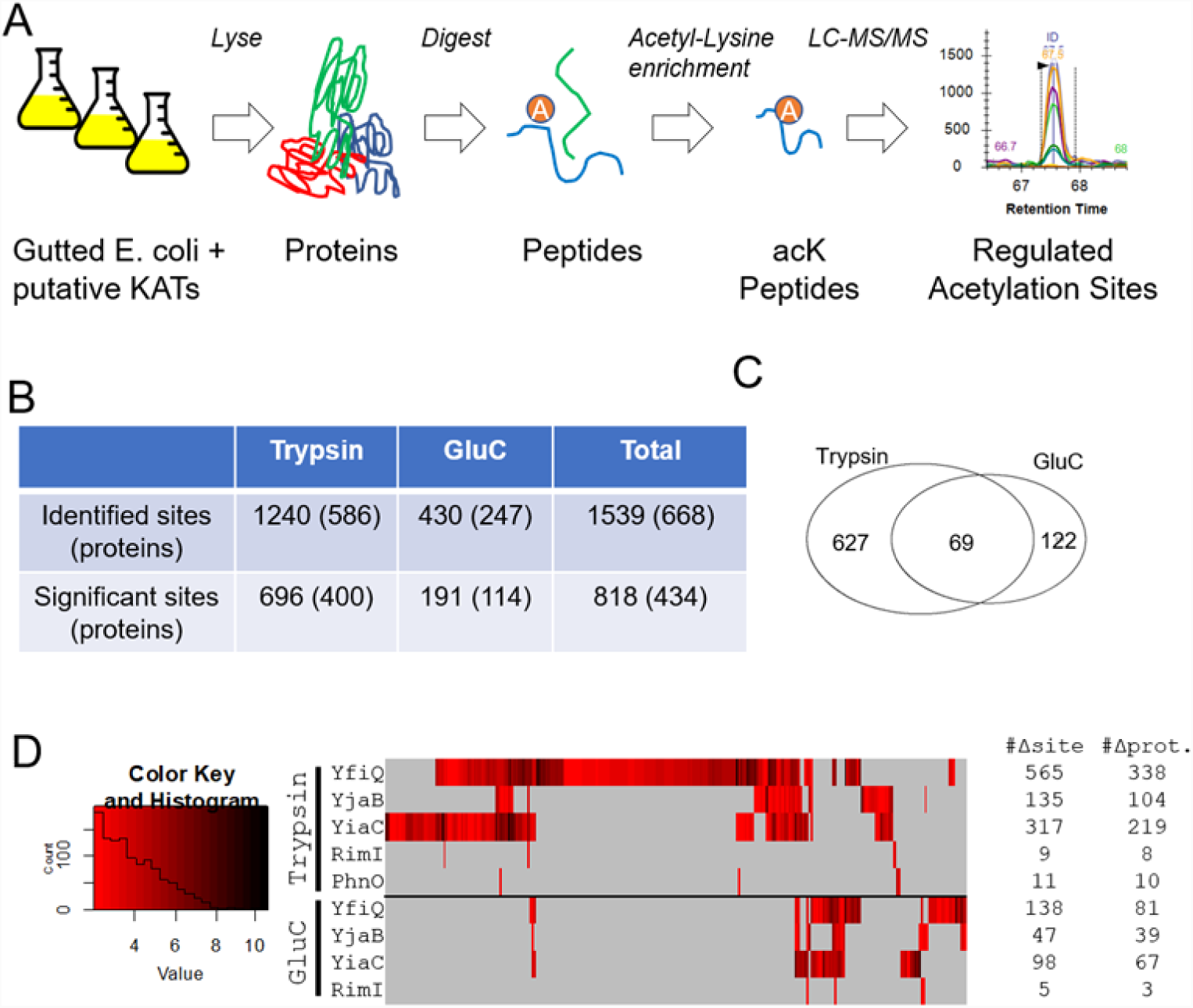
Identification of site-specific regulation of acetylation sites by KATs. (A) Cartoon showing workflow used to identify KAT target sites. (B) Overview of the significant sites and proteins regulated by at least one putative KAT. Significance defined as FDR < 0.01 within-set and log2(FC) > 2. (C) Venn Diagram showing the complementary nature of trypsin and GluC digestion in terms of significant acetylation sites. (D) Heat map of all significant changes; acetylation sites are grouped by unsupervised hierarchical clustering.

To further explore the specificity of these KATs, we compared the sites acetylated by KAT overexpression with sites that we previously found to be sensitive to deletion of *ackA*, which causes accumulation of the highly reactive acetyl donor AcP and therefore results in non-enzymatic protein acetylation (**Fig. S5A, Table S1C**) (5). Remarkably, of the 592 ackA-regulated sites, only 29 overlapped with the 818 sites acetylated by PhnO, RimI, YiaC, YjaB, or YfiQ, further reinforcing their specificity and thus likely distinct functions. We also analyzed the primary amino acid sequences surrounding lysines that were acetylated by these novel KATs and found no specific neighboring residue preference (**Fig. S5B, S5C**). This suggests substrate specificity cannot be determined by primary sequence alone and that three-dimensional analysis of protein structures should be taken into account.

### KAT-dependent acetylation of proteins involved in translation and glycolysis

A large number of KAT substrate proteins were found to be involved in the GO Biological Process term translation (Benjamini-corrected p-value 1.1E-22, determined by DAVID functional enrichment tool) (28, 29). Almost all ribosomal protein subunits were acetylated (51 of 55 proteins); some were acetylated by AcP only (8/55), some were acetylated by one or more KATs but not AcP (11/55), but most were acetylated by at least one KAT and AcP (32/55) (**Table S2A**). Very few ribosomal lysines were acetylated both by a KAT and AcP (only 9 of 184 sites on the 55 subunits). In contrast, one-quarter (46/184) of the observed acetylated lysines were targeted by more than one KAT, with as many as 3 KATs acetylating the same lysine. Most of the amino acid-tRNA ligases (16/23 proteins) were acetylated; some were acetylated by AcP only (6/23), some by KATs only (5/23) and some by both (5/23). Again, lysines that were acetylated by both a KAT and AcP were rare (2/41 sites on 23 proteins). Only a few lysines were acetylated by multiple KATs (4/41). Three of the 7 elongation factors were acetylated; these acetylations were largely dependent on AcP (13/15). All of the initiation factors were acetylated, and these acetylations were almost entirely KAT-dependent (7/8). These results are consistent with distinct roles for KAT-dependent and AcP-dependent acetylations.

Regarding the interplay between non-enzymatic and enzymatic acetylation, central metabolism was particularly interesting (**Fig. 5**). Twenty-seven proteins comprise the 3 glycolytic pathways in *E. coli* (Embden-Meyerhof-Parnas [EMP], Entner-Dourdoroff [ED], and Pentose Phosphate [PP]). Of these 27 proteins, 20 were detected as acetylated: 2 strictly by KAT(s), 7 by KAT(s) and AcP, and 11 by AcP alone. A total of 97 lysines were acetylated: 9 by KAT(s) alone, 86 by AcP alone, and only 2 by both AcP and a KAT. Intriguingly, the majority of KAT-dependent acetylations (7/11) were found on proteins responsible for either the early or late steps of glycolysis, i.e. prior to the formation of glyceraldehyde 3-phosphate (GAP) or on enzymes responsible for aerobic AcCoA synthesis. In contrast, the majority of AcP-dependent acetylations (66/88) were found on proteins that all glycolytic pathways share. In support of the concept that KAT-dependent acetylation helps direct flux, 3 other proteins relevant to glycolysis are exclusively acetylated by KAT(s). YfiQ and YiaC acetylated the transcription factor GntR, which controls expression of the enzymes (Eda [2-keto-4-hydroxyglutarate aldolase] and Edd [phosphogluconate dehydratase]) that comprise the ED pathway (30). LipA synthesizes lipoate, whereas LipB transfers a lipoyl group onto a lysine in the E2 subunit (AceF) of the pyruvate dehydrogenase complex (PDHC). The 3 subunits of PDHC, whose activity requires lipoylation, are highly acetylated, but almost entirely by AcP. In contrast, LipA and LipB are entirely acetylated by KATs (7 lysines on LipA by YfiQ, YiaC, and YjaB and 1 lysine on LipB by YfiQ). These observations are consistent with the hypothesis that KAT-dependent acetylation helps direct flux through the 3 different glycolytic pathways and regulates the transition from glycolysis to AcCoA-dependent pathways, such as the TCA cycle, fatty acid biosynthesis, and different forms of fermentation.

**FIGURE 5.**
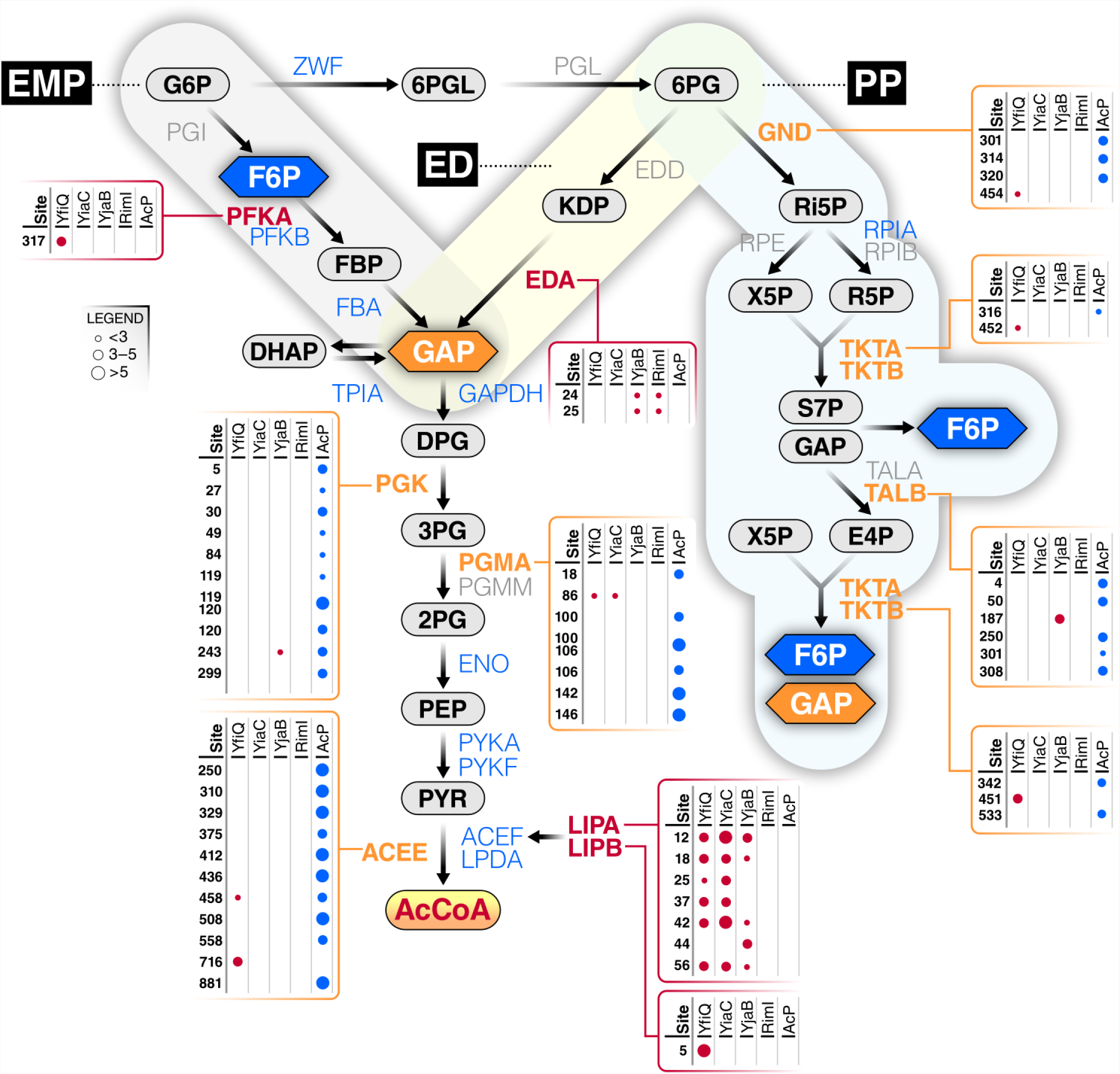
Most of central metabolism is differentially acetylated by acetyl-P and/or KATs. The three glycolytic pathways, Embden-Parnas-Meyerhof (EMP), Entner-Dourdoroff (ED), and Pentose Phosphate (PP) are shown with metabolites and enzymes indicated. Some enzymes are not acetylated (gray), while others are acetylated by acetyl-P alone (blue), KATs alone (red), or both (orange). Enzymes with boxes were modified by at least one KAT (as indicated); some were also acetylated by acetyl-P (AcP). The size of the dot indicates the fold upregulation for each lysine by either a KAT or AcP.

### Structural analysis of KAT and AcP-dependent acetylation sites

Previously, we analyzed the location of lysine residues on several glycolytic enzymes that are non-enzymatically acetylated by AcP (5). Here, we expanded our structural analysis to include selected enzymes in the EMP, ED, and PP pathways. We specifically investigated the KAT-dependent and/or AcP-dependent acetylated lysines on available *E. coli* protein structures (**Fig. 6**). Excluding proteins modified by AcP alone, we evaluated acetylated proteins from three main groups: acetylated by a KAT only, acetylated by a KAT and AcP on different lysines of the same protein, and acetylated by a KAT and AcP on the same lysine of the same protein. Examples of proteins that were only acetylated by a KAT included PfkA and Eda, those modified by either a KAT or AcP on different residues included PgmA and TalB, and those modified by a KAT and AcP on the same lysine residue included Pgk. One representative protein from these pathways that was modified by each individual KAT was selected to evaluate substrate lysine locations in 3D (**Fig. 5; Fig. 6 A-D**): PfkA (YfiQ; EMP), Eda (RimI; ED), PgmA (YiaC; EMP), and TalB (YjaB; PP). Note that some of these proteins are modified by multiple KATs, a scenario that we will discuss later.

**FIGURE 6.**
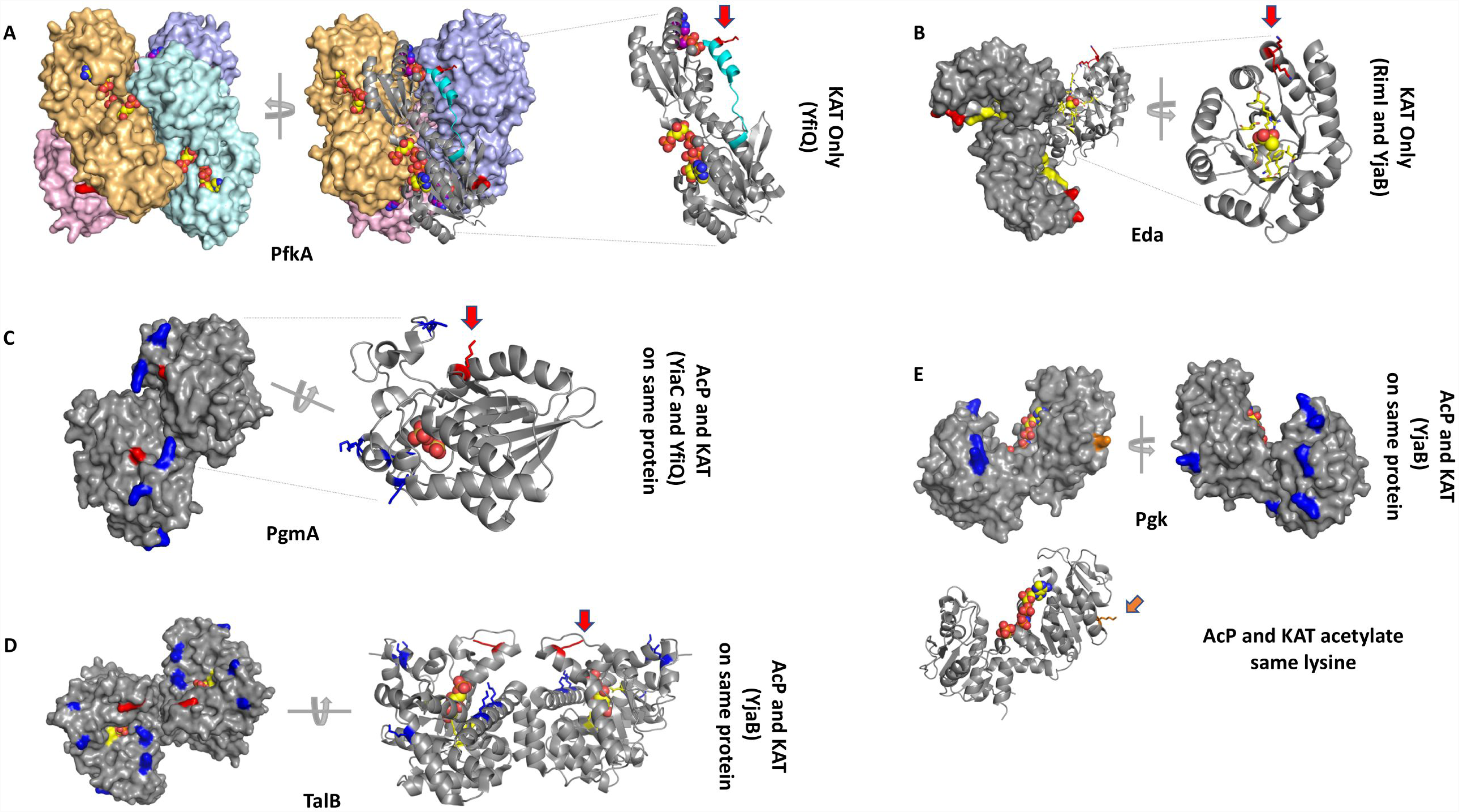
Structural analysis of selected proteins modified by KATs and/or AcP. (A) PfkA (PDB ID: 1pfk) structure. Each monomer of the tetramer is tinted in pink, orange, cyan, and violet, and shown as a surface representation. The ligand ADP is bound to the allosteric site and the ligand fructose 1,6-bisphosphate is bound to the active site; both are shown as spheres. One monomer is also shown as a ribbon representation. A red arrow indicates the location of K317. The C-terminus that is disordered in the 2pfk structure is shown in cyan. (B) Eda (PDB ID: 1eua) structure. A surface representation of the trimer is shown in gray. One monomer of the trimer is also shown as a ribbon representation and a red arrow indicates the two adjacent sites of acetylation. Pyruvate is shown as spheres. The active site residues are colored in yellow and K24 and K25 are colored in red. (C) PgmA (PDB ID: 1e58) structure. The dimer is shown as a surface representation and one monomer of the dimer is shown as a ribbon representation. Sulfate is shown as spheres in the active site. K86, which is acetylated by YfiQ and YiaC is shown in red and indicated by a red arrow. K18, 100, 106, 142, and 146 are acetylated by AcP and shown in blue. (D) TalB (PDB ID: 4s2c) structure. The dimer is shown in both surface and ribbon representations. K187 is acetylated by YjaB and shown in red with a red arrow. K4, 50, 250, 301, and 308 are acetylated by AcP and shown in blue. Fructose 6-phosphate is shown as spheres in the active site, and surrounding residues are shown as yellow sticks. (E) Pgk (PDB ID: 1zmr) structure. The monomer is shown as a surface and ribbon representation where K243, which is acetylated by both YjaB and AcP, is shown in orange and an orange arrow points to its location. K5, 27, 30, 49, 84, 119, 120, and 299 are acetylated by AcP and shown in blue. Phosphoaminophosphonic acid-adenylate ester and 3-phosphoglycerate are shown as spheres in the active site and modeled from the 1vpe structure.

#### Comparison of KAT only acetylated lysines on selected substrate proteins

Phosphofructokinase A (PfkA) is an allosterically regulated tetrameric protein. We found that the acetylated lysine (K317) of PfkA was located at the end of an α helix at the C-terminus of the protein and lays in a pocket formed by a second monomer of the tetramer (**Fig. 6A**). Therefore, K317 is found at the interface between monomers of the tetramer and lies outside the active site and allosteric site of the protein. K317 forms a salt bridge with D273 of an adjacent monomer, and Paricharttanakul (31) previously found that D273 is likely important for stabilizing the tetramer and affects the allosteric activation and inhibition network. A disruption of this salt bridge via acetylation could possibly alter allosteric properties of the protein. Furthermore, the C-terminus is important for stability of the oligomer (32), and when the allosteric effector ADP is not bound, this region becomes disordered (33). The fact that this region is disordered in the absence of the allosteric effector in the crystal structure suggests this portion of the protein is mobile and therefore may be accessible to YfiQ for acetylation.

KHG/KDPG aldolase (Eda) is a trimeric protein and was found to be acetylated on two adjacent lysines (K24, K25) by both RimI and YjaB (**Fig. 6B**). These amino acids are found at the end of a surface accessible α helix, which is outside of the active site and is not at the interface of monomers of the trimer. The only interaction observed for either of these amino acids was a salt bridge between K25 and E193 on a neighboring α helix. For this reason, the function of lysine acetylation on this protein is unclear.

#### Comparison of KAT and AcP acetylation sites on different lysines of the same protein

Phosphoglycerate mutase (PgmA) contains six lysines that were acetylated by either AcP or a KAT (**Fig. 6C**). K86 was the only site of enzymatic acetylation (YiaC and YfiQ), whereas K18, 100, 106, 142, and 146 were all non-enzymatically acetylated by AcP. While this enzyme was not previously considered to be allosteric, it was recently proposed to function as an allosteric enzyme, whereby dimer stabilization acts as the allosteric signal that is transmitted rather than the more typical binding of a specific effector to an allosteric site. In this case, the ordering and stabilization of the region that contains the lysines acetylated by AcP acts as the transmission signal (34, 35). While all AcP acetylations occurred on this highly flexible domain of the protein, the KAT acetylation site (K86) was found outside the active site on a small 3_10_-helix near but not directly interacting with the opposite monomer at the dimer interface. This lysine coordinates a water molecule between itself and E166 on a neighboring α helix. However, if stabilization of the dimer is truly acting as the allosteric signal, then this lysine (K86) is only indirectly involved in the allosteric site. Investigation of the AcP modified lysines showed that only K100 was found to be in the active site. The function of all other lysines that were acetylated by AcP are currently unclear.

Transaldolase B (TalB) is a dimeric protein that is modified through both enzymatic and non-enzymatic acetylation (**Fig. 6D**). Similar to PgmA, TalB is also acetylated on one lysine by a KAT (YjaB; K187) and several lysines by AcP (K4, 50, 250, 301, and 308). All these acetylated lysines are found on α helices on the same face of the C-terminal side of the beta-barrel. The α helices that surround the beta-barrel are known to be mobile (36). The enzymatic acetylation site occurs near the end of an α helix outside of the protein active site and is not found at the interface between monomers of the dimer. There appears to be some specificity of acetylation in this location of the protein because two additional lysines are directly downstream (K192 and 193) on a loop and are not acetylated by either a KAT or AcP. Lysines acetylated non-enzymatically by AcP are also surface accessible. Two of these lysines (K301 and 308) are found on a long α helix that creates the dimer interface, but neither participates in interfacial interactions. K301 is near the active site where sugar phosphates bind, but K308 is further down the helix. This C-terminal helix is preceded by an extremely long loop (residues 254-277) that connects it with two additional helices that contain AcP-modified lysines K4 and K250. K4 forms polar interactions between its ε amino group and the backbone carbonyl oxygens of both S255 and E256, while K250 forms a salt bridge with E254. The ε amino group of K50 also forms polar contacts with the backbone carbonyl oxygen of E46, but is not linked to the long loop or helix where other AcP-modified lysines are found. It is unclear what effect these acetylated lysines have on protein function or oligomerization.

#### KAT and AcP acetylation of the same lysine residue

Only select lysines within each substrate protein are acetylated. Most often, the method by which these lysines become acetylated is either exclusively by a KAT or by AcP. In rare cases, both mechanisms compete for the same lysine. One example where this occurs is in phosphoglycerate kinase (Pgk), whereby both AcP and a KAT (YjaB) acetylate K243 (**Fig. 6E**). Pgk is monomeric and the *E. coli* protein has been crystallized in the open conformation. In other homologs, the structures of the partially closed and fully closed forms of the protein have also been determined (37). K243 is found at the end of an α helix and has no specific interactions with other amino acids of the protein. AcP also acetylated lysines 5, 27, 30, 49, 84, 119, 120, and 299. All KAT and/or AcP acetylated lysines are found on the surface of the protein and do not interact directly with the active site. Two lysines acetylated by AcP (K27 and 30) are on a loop that moves upon closure of the protein. Nearly all lysines acetylated by AcP are found within the N-terminal domain, whereas K243 and K299 are located in the C-terminal domain. It is unclear why both KAT and AcP acetylate K243, and why the other lysines are preferred sites for acetylation by AcP.

### Structural and active site residue comparison of KATs

Initially, we used Phyre2 to predict amino acids that may be involved in catalysis of KATs, and these predictions informed our mutagenesis trials. Here, we chose to perform a more thorough structural analysis to determine whether these suggested amino acids were present in locations known to be important for activity in homologs. The sequence identity between KATs is low (<30%), but since GNATs share the same structural fold, we performed a structural comparison of these KATs in order to identify the location of active site residues in 3D. The *E. coli* crystal structure of RimI (5isv) and NMR structure of YjaB (2kcw) have been deposited into the Protein Data Bank (PDB); however, no structures have been determined for the other *E. coli* KATs (YfiQ, YiaC, and PhnO). Therefore, we built homology models of these three proteins.

Based on our models and available structures, we found that all KATs adopted the standard GNAT fold with a characteristic V-like splay. The structures and models also informed our manual refinement of our sequence alignment of KATs (**Fig. 7A & B**). There is significant sequence and structural variability in the α1-α2 and β6-β7 regions of each KAT. However, all of their active sites, with the exception of YfiQ, contained a conserved tyrosine known to act as a general acid in other GNAT homologs (21, 38). Upon further analysis, we found that the identity or location of the amino acid that tends to act as a general base may not be as conserved across KATs compared to the amino acid that acts as the general acid. For instance, E103 coordinates a water molecule to act as a general base in RimI from *Salmonella typhimurium* LT2 (21) is in the same location in 3D as the corresponding amino acids (both E103) in *E. coli* RimI and YiaC (**Fig. 7C**). In contrast, N105 and S116 are in the same location in YjaB and PhnO, respectively. In theory, these amino acids can coordinate a water molecule, but to our knowledge the effect of substituting these amino acids for glutamate has not been evaluated. Our mutagenesis of E78 in PhnO significantly decreased its acetylation activity in the “gutted” strain (**Fig. 3**), indicating that this amino acid is critical for catalysis. Thus, the location of the amino acid that either coordinates a water molecule that functions as the general base in the reaction or the amino acid that directly participates in this function may be in a different location in 3D on different KATs. Mutation of F70 in YiaC decreased acetylation activity (**Fig. 3**), but not as substantially as the tyrosine mutation. In 3D, the equivalent amino acid in all KATs is hydrophobic (**Fig. 7C**), which may be important for substrate recognition. Regardless, this amino acid does not directly participate in the chemical reaction.

**FIGURE 7.**
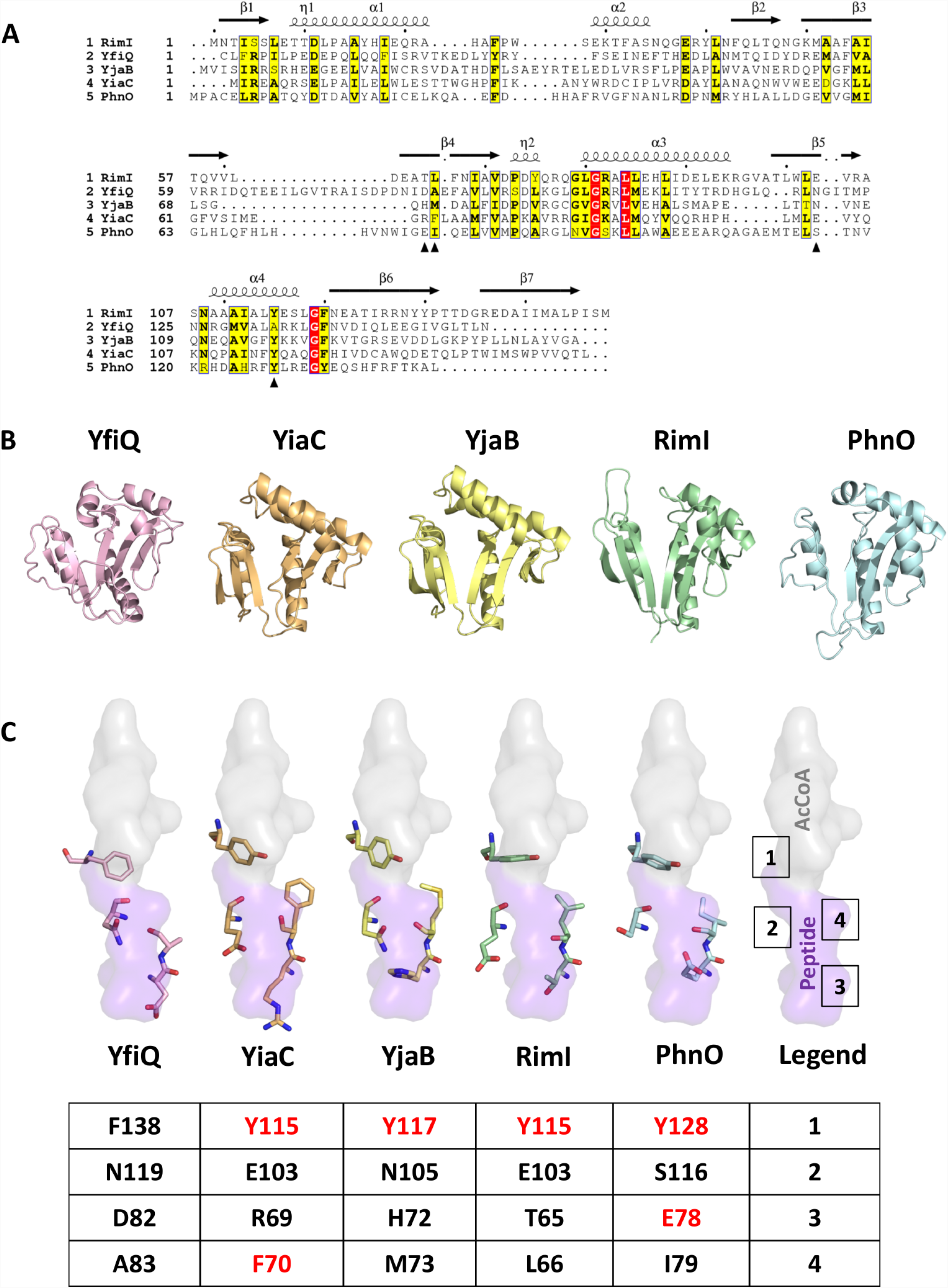
Sequence and structural comparison of KAT proteins and their key catalytic residues. (A) Sequence alignment of all five *E. coli* KATs. Only the GNAT portion of the sequence for YfiQ is shown. The structural elements above the sequences are based on the 5isv RimI structure. Red highlighting represents 100% identity, whereas yellow highlighting shows a global score of 70% identity based on ESPript 3.0 parameters. Black arrows beneath the sequences indicate the residues selected for structural comparison in panel C. (B) Comparison of overall structures and homology models of *E. coli* YfiQ (pink), YiaC (orange), YjaB (yellow), RimI (green; full C-terminus not shown in the figure), and PhnO (blue) proteins in a ribbon representation. 3D structures of YjaB and RimI were determined previously (PDB IDs 2kcw and 5isv, respectively). We built homology models of the remaining KATs using the following structures as templates: 4nxy for YfiQ, 2kcw for YiaC, and 1z4e for PhnO. Only the GNAT portion of the YfiQ protein sequence was used for the homology model. Further details regarding parameters for building and selecting representative homology models for these proteins are described in Materials and Methods. (C) Comparison of select active site residues potentially important for substrate recognition and catalysis in GNATs. The crystal structure of RimI (5isv) has the C-terminus of one monomer bound in the active site of the second monomer. A surface representation of this portion of the protein that encompasses the AcCoA donor (gray) and peptide acceptor (purple) site are shown. Each of the KAT homology models and structures were aligned using TopMatch and Pymol. Four active site residues are shown. A table beneath the structures shows the specific residue numbers for each KAT. Residues that were mutated are shown in red.

### Newly identified KATs are conserved

Having identified these KATs in *E. coli*, we next asked whether orthologs of these KATs were present across bacterial phylogeny. To find these orthologs, a Hidden Markov Model (HMM) was built for each gene of interest and searched against 5589 representative reference genomes from RefSeq (39, 40). Cutoffs for the HMM match score were derived manually by identifying a score such that genomes contained at most only one match. This cutoff was chosen to draw the line between orthologs and paralogs, i.e. when a genome has multiple copies of similar sequences but only one contains the biological function of the query sequence. As a highly conserved gene, *rimI* was identified in 4459 genomes (**Table S3A**). The *yiaC* and *yjaB* genes were all broadly distributed across bacterial taxa and found in 421 and 692 genomes, respectively (**Table S3B**). However, *phnO* was found to have a very limited distribution, identified in only 22 genomes and appears to belong exclusively to the γ-proteobacteria. A representative phylogenetic tree shows the broad distribution of *yjaB* across the bacterial domain (**Fig. 8**).

**FIGURE 8.**
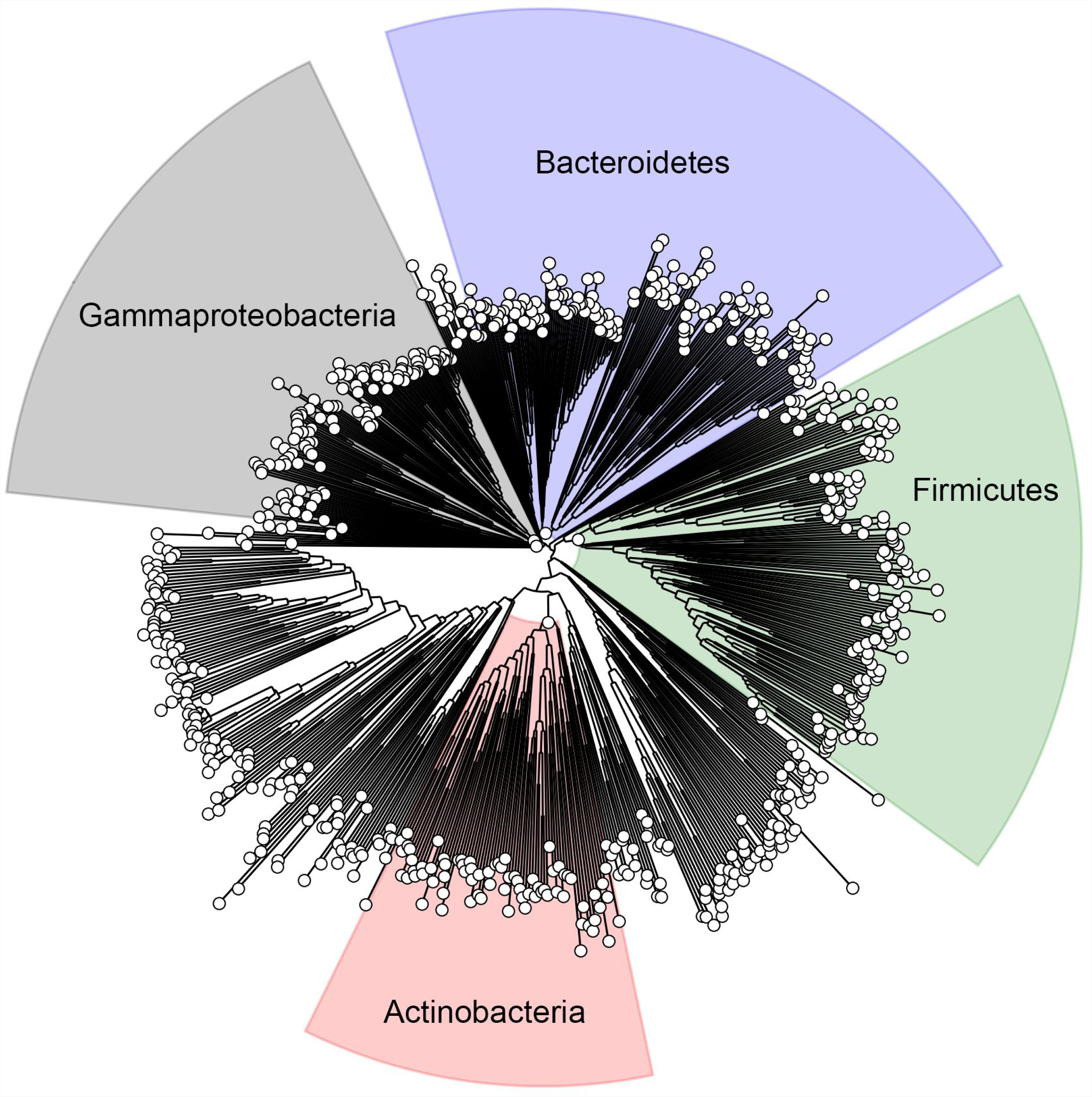
YjaB is highly conserved across bacteria. A representative phylogenetic tree showing bacterial species that contain a gene homologous to *yjaB* from *E. coli*. The trees for *rimI* and *yiaC* are comparable.

19 genomes contained all four new KATs. Unsurprisingly, many of these genomes correspond to *E. coli* strains or the closely related species *Salmonella enterica*. 148 genomes contained three KAT and 782 genomes contained two KATs. Study of these KATs expressed heterologously in *E. coli* could help to understand the potential role of acetylation in these other bacteria.

### YiaC and YfiQ can inhibit migration in soft agar

We sought to determine the physiological relevance of acetylation by these KATs. Based on the *E. coli* gene expression database (https://genexpdb.okstate.edu/databases/genexpdb/), we found conditions when these KATs may be expressed and, thus, when they may be relevant. Expression of each of the KATs appeared to be upregulated in stationary phase and/or biofilm conditions. Thus, we tested overexpression constructs of the four novel KATs and YfiQ in a mucoidy assay, but we did not observe any difference relative to wild-type cells. We then tested these strains for motility. We found that overexpression of YiaC and YfiQ consistently reduced migration in a soft agar motility assay (**Fig. 9A**). The inhibition of migration was not due to a reduction in growth rate as the overexpression strains grew as well as their vector controls (data not shown). To determine whether this reduction required the acetyltransferase activity of YiaC, we tested overexpression of YiaC F70A, which had reduced acetyltransferase activity, and YiaC Y115A, which lost activity (**Fig. 9B**). Overexpression of YiaC YF70A inhibited migration similar to overexpression of wild-type YiaC. In contrast, YiaC Y115A was unable to inhibit migration. To ensure this was not a strain specific phenomenon, we recapitulated these data for YiaC in another strain background, MG1655 (**Fig. S6**). However, the overexpression of YfiQ caused a growth inhibition in MG1655 (**Fig. S6**). If YiaC and YfiQ inhibit motility, then deletion of those genes may increase migration. However, the Δ*yiaC* mutant migrated equivalently to the wild-type parent, while the Δ*yfiQ* mutant had a slight reduction in migration in BW25113 (**Fig. 9D**).

**FIGURE 9.**
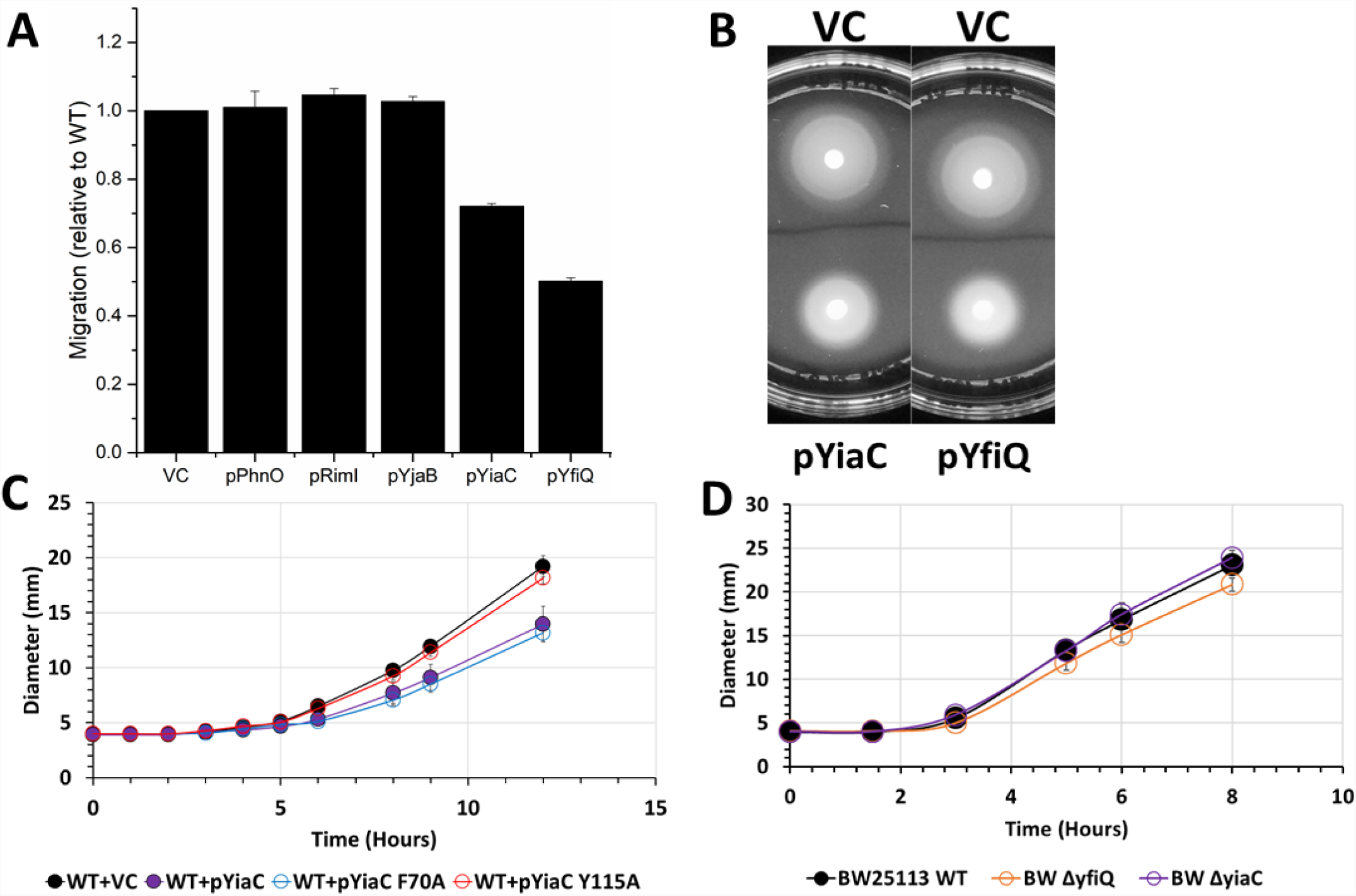
YiaC and YfiQ inhibit migration. Cultures were grown overnight in TB medium supplemented with chloramphenicol and 50 μM IPTG. 5 μL of each normalized culture was spotted on low percentage TB plates supplemented with chloramphenicol and 50 μM IPTG. The diameter of the cell spot was measured hourly. (A) Final diameter relative to vector control (VC) after 12 hours is shown for wild-type *E. coli* strain BW25113 strains carrying the indicated plasmids. (B) Representative motility plates of BW25113 carrying VC, pYiaC, or pYfiQ. (C) Hourly migration of wild-type *E. coli* strain BW25113 strains carrying pCA24n encoding YiaC, YiaC mutants, or vector control (VC). (D) Hourly migration of wild-type *E. coli* strain BW25113 or isogenic deletion mutants on low percentage TB plates without supplement.

## DISCUSSION

Over the last decade, Nε-lysine acetylation has become recognized as an important post-translational modification of bacterial proteins that regulates physiology. While acetylation of certain lysines may have a clear output, such as inhibition of enzyme activity due to active site Nε-lysine acetylation, the functional importance of many acetyllysine modifications are more difficult to discern. To uncover the role of acetylation in these unclear cases, it is helpful to use a model bacterium (e.g., *E. coli*) with a vast knowledgebase of pathways, protein structure-function relationships, and physiology. Therefore, we constructed a “gutted” strain that lacked both known acetylation mechanisms (5, 6, 23) to examine if KATs other than YfiQ exist. This approach substantially decreased background acetylation, increased the signal-to-noise ratio, and allowed us to identify four enzymes that possess robust KAT activity: RimI, YiaC, YjaB, and PhnO. We acknowledge that overexpression may produce artifacts. However, knowledge of amino acids required for catalytic activity in homologous enzymes allowed us to construct inactive or minimally active mutant enzymes and determine that they function as KATs.

To define statistically significant KAT lysine target sites, we applied very stringent requirements of >4-fold increase in acetylated lysines in the KAT overexpression strains relative to the vector control and a q-value of less than 0.01. Using these strict criteria, we identified 818 acetylated lysines on 434 proteins. Most of these modified lysines were acetylated by a single KAT. While the overlap of lysines acetylated by the 5 KATs is relatively minor, the overlap between all the KAT-dependent acetylations and AcP-dependent acetylations is even smaller. The specificity of each KAT suggests that *E. coli* has evolved distinct regulatory modalities, perhaps reflecting the need to remodel the proteome in certain environments. This concept is supported by the patterns of acetylation of translation-associated proteins and of the glycolytic pathway proteins (**Fig. 5**). To further emphasize this concept, we determined that YfiQ and YiaC, but not the other three KATs, inhibit motility. This suggests that acetylations catalyzed by YfiQ and YiaC have distinct outcomes and thus are specific. Determining conditions when these KATs are active could reveal the advantage of minimal redundancy among targets.

The possibility that GNAT family members other than YfiQ might possess KAT activity was examined in two studies by Venkat and co-workers, where the ability of GNATs to *in vitro* acetylate malate dehydrogenase and tyrosyl-tRNA synthetase was assessed (41, 42). Neither protein was enzymatically acetylated by GNATs, at least not above the acetylation level achieved by incubating with AcCoA alone. Importantly, our mass spectrometry data corroborate those data, as neither malate dehydrogenase nor tyrosyl-tRNA synthetase were modified by the KATs we identified *in vivo*. With similar *in vitro* acetylation assays performed by our group, the level of acetylation is already high on purified target proteins, so the effect of the KATs on acetylation of the targets was small or unobservable. We are currently working on optimizing this protocol to investigate the activity of these KATs *in vitro*.

Out of the hundreds of proteins we identified as acetylated by the newly identified KATs, several are central metabolic proteins and many are components of the translational machinery. From our structural analysis of a limited set of proteins, it appears that KAT-dependent acetylations occur primarily on the ends of α helices near allosteric or active sites, and sometimes at oligomeric interfaces. Most sites tend to be surface accessible and may be intricately involved in allosteric signaling networks and/or mediate protein-protein interactions. On the other hand, AcP-dependent acetylations are mainly located on α helices, loops, and active site amino acids. Additional studies are needed to understand why multiple KATs acetylate the same lysine on the same protein. Future studies will decipher the role of specific target proteins, such as LipA, that are differentially acetylated by multiple different KATs and may be differentially regulated depending on stress, environment and nutrient availability. Similarly, it will be important to determine if KAT-acetylated lysines on helices have the propensity to unwind compared to other helices in the KAT substrate proteins or to helices containing AcP-acetylated lysines. To examine whether these trends hold for a larger set of KAT substrate proteins, we are currently performing a wider analysis across all identified substrates with structures.

At this point, we can only speculate on the effects these modifications have on many of the target proteins, but for some proteins not included in our structural analysis, critical lysines are acetylated. For example, selenide water dikinase (also known as selenophosphate synthetase) is acetylated by YiaC on K20, an amino acid known to be critical for catalyzing selenophosphate synthesis (43). Both YiaC and YfiQ acetylated FabI on K205, an amino acid known to be important for an essential step in fatty acid biosynthesis: the reduction of an enoyl-acyl carrier protein (44). YfiQ acetylated adenylate kinase (Adk) on K136, which is known to be important in stabilizing the open state of the enzyme by forming a salt bridge with D118. This salt-bridge appears to be important for dynamic transitions between different states (45). Finally, cysteine synthase A (CysK) is acetylated by YiaC on K42, which is within the active site of the protein and becomes covalently modified by pyridoxal 5’-phosphate (PLP); the enzymatic activity of CysK depends on Schiff base formation of this lysine with PLP (46). In the case of large protein complexes, such as the ribosome, that are multiply acetylated on several subunits, it is tempting to speculate that acetylation of some seemingly inconsequential lysines may produce significant effects when combined; for example, stabilizing or destabilizing complex formation or altering ribosomal function.

We find proteins in pathways involved in metabolism and translation are particularly heavily acetylated. However, many KAT-dependent acetylations in these processes were distinct from those catalyzed by AcP as previously reported (5, 6) The KATs described here tend to acetylate enzymes that regulate the branch points of central metabolism, while AcP seems to modify many of the central metabolic enzymes (**Fig. 5**). This suggests the tempting hypothesis that KATs have evolved to specifically regulate key flux points in metabolism, while AcP-dependent acetylation may be a global response to the carbon flux.

A recent report revealed the only known *E. coli* deacetylase CobB has lipoamidase (delipoylase) activity (47). Lipoyl groups can be found on subunits of several major central metabolic complexes and contribute to the activity of these complexes. Rowland *et al.* found that CobB could regulate the activities of the pyruvate dehydrogenase (PDH) and α-ketoglutarate dehydrogenase (KDH) complexes and could delipoylate the AceF and SucB components of PDH and KDH, respectively. As mentioned in the results above, we find that most of these metabolic complexes are multiply acetylated by AcP and/or KATs, including AceF and SucB. Additionally, we found that a protein responsible for generating the lipoyl groups for these lipoylated subunits, LipA, was highly acetylated by KATs, and Rowland *et al.* found that LipA co-immunoprecipitated with CobB. The tight co-occurrence of lipoylated and acetylated proteins at key nodes of central metabolism with the potential to be regulated by CobB suggests an interesting dynamic between acetylation and lipoylation that warrants further study.

*E. coli* RimI appears to acetylate lysines on multiple proteins. This is an interesting observation, as RimI from both *E. coli* and *Salmonella typhimurium* is known to function as an N-terminal alanine acetyltransferase that has but one known target, the ribosomal protein S18 (21, 48, 49). While RimI from *E. coli* and *S. typhimurium* is characterized by stringent N-terminal alanine specificity, RimI from *Mycobacterium tuberculosis* exhibits relaxed N-α amino acid substrate specificity *in vitro* (50). Intriguingly, we observed that *E. coli* RimI can also acetylate an Nε-lysine on a different ribosomal protein, L31. The Nε-lysine of L31 is found on a long unstructured region of the protein and may bind to RimI in a similar conformation as the C-terminus of the *E. coli* RimI protein in its crystal structure (PDB ID: 5isv). The α-amino group of alanine on S18 is also found at the end of a long unstructured region. This insinuates that RimI could exhibit both Nα-amino acid and Nε-lysine acetylation activity, but the substrate specificity of this enzyme is still unclear.

Similarly, PhnO is an aminoalkylphosphonate acetyltransferase in both *E. coli* and *S. enterica* (51, 52). PhnO is part of a gene cluster involved in the utilization of phosphonate under inorganic phosphate starvation conditions. While PhnO is not absolutely required for phosphonic acid utilization, it does acetylate (S)-1-aminoethylphosphonate and aminomethylphosphonate (52, 53). Of the 10 proteins PhnO acetylates, one is inorganic triphosphatase, which indicates additional levels of phosphate regulation via this KAT.

Prior to our study, both RimI and PhnO were identified as having functions unrelated to Nε-lysine acetylation. It is interesting to note that these two KATs have significantly fewer internal lysine protein substrates (11 and 10, respectively) than do YfiQ, YiaC, or YjaB. The reasons for this dual character of the two KATs are unknown. While it could simply be that these enzymes have broad substrate specificity, it is tempting to speculate that the Nε-lysine acetylation by RimI and PhnO may be part of a more complex cellular regulatory mechanism for bacteria that harbor these KATs.

We found that overexpression of both YiaC and YfiQ inhibit motility. For YiaC, none of the targets that we determined by mass spectrometry provide a simple explanation for this phenotype. For YfiQ, the effect of deletion or overexpression on motility has not been directly assessed, although a previous report supports the idea that YfiQ could inhibit motility, as a Δ*yfiQ* mutant exhibited slightly enhanced transcription of flagellar genes (54). However, there is also evidence that YfiQ can enhance motility. This report and others find that YfiQ can acetylate K180 of RcsB, a response regulator that represses transcription of the master regulator of flagellar biosynthesis, *flhDC*. Acetylation of K180 would thus be expected to prevent repression by inhibiting RcsB from binding to DNA, enhancing migration (55). This is contrary to what we observe, which suggests that YfiQ inhibits migration through a target other RcsB, at least under the tested conditions. Finally, YfiQ acetylates K67 and K76 of FlgM, an anti-sigma factor for the sigma factor FliA (σ^28^). FliA is required for initiation of many genes involved in flagellar biosynthesis. If YfiQ was acting through FlgM to inhibit motility, it would suggest that acetylated FlgM would bind FliA more tightly.

Contrary to our expectations, both Δ*yfiQ* and Δ*yiaC* mutants migrated at a rate similar to their wild-type parent. There are three possible explanations: (1) YfiQ and YiaC do not regulate motility, (2) YfiQ and YiaC may not be expressed under the tested conditions and thus deletion of these genes would have no effect and (3) YfiQ and YiaC may compensate for each other; alternatively, some other KAT or AcP may contribute to compensation.

While we have a phenotype for overexpression of YiaC, the other three new KATs do not yet have clear phenotypes. Analysis of the *E. coli* gene expression database suggested conditions under which these KATs may be expressed. Based on these data, RimI (56-58), PhnO (59, 60), YjaB (61, 62), and YiaC are all upregulated during stationary phase dependent on the stationary phase sigma factor, RpoS, and in biofilm-forming conditions. RimI (58), PhnO, and YjaB (62, 63) are upregulated in heat shock conditions, while PhnO and YiaC are downregulated during cold shock. Furthermore, YiaC is upregulated during oxidative stress (63). We also searched the genomic context and found YiaC and PhnO are encoded in polycistronic operons, while YjgM and YjaB are monocistronic. The *yiaC* gene is directly downstream and overlaps four nucleotides of the *tag* gene that encodes 3-methyl-adenine DNA glycosylase I. The product of the *tag* gene is important for removing potentially mutagenic alkylation damage from DNA, but it is not induced through the adaptive response. This may suggest a separate promoter for *yiaC* within the *tag* gene that allows it to respond to oxidative stress. As mentioned previously, the *phnO* gene is encoded with the other genes necessary for phosphonate utilization. We are currently pursuing phenotypic analyses of these KATs based on this information and the targets that we have identified.

Excitingly, these KATs are well conserved across bacteria. Thus, discoveries about how these KATs affect *E. coli* physiology are likely applicable to other bacteria. For example, many organisms require motility for their pathogenicity, and our data suggests that YiaC and YfiQ may regulate motility in *E. coli*. However, both *Yersinia pestis* and *Klebsiella pneumoniae* encode YiaC and the fact that both species are non-motile suggests other roles for these enzymes in these particular bacteria. KAT homologs are also encoded in pathogens such as *Listeria monocytogenes* and *Pseudomonas aeruginosa*, and it would be interesting to determine whether these KATs regulate pathogenesis. A simple method to determine whether these homologs possess KAT activity would be to use our “gutted” approach. By expressing a heterologous putative KAT in our “gutted” strain, one could perform western blot or mass spectrometry to determine any changes to the acetylome. Indeed, we have evidence that this can work for at least one protein from *Neisseria gonorrhea* (Christensen *et al*., unpublished data). However, it is important to note that *E. coli* may not encode the targets from the native species, so determination of native targets must be validated independently.

In conclusion, we identified four GNAT family members that have KAT activity in addition to the known KAT, YfiQ. These five KATs catalyze acetylation of hundreds of proteins on over 1500 lysines, and the acetyltransferase activity depends on conserved catalytic tyrosines and/or key glutamates found in many GNAT family members. Furthermore, the conservation of YiaC in certain pathogenic organisms like *Yersinia pestis* warrants consideration as a topic of study. Clearly, our results provide a starting point for further analysis that is sure to yield fruitful mechanistic and regulatory insight into the complex orchestration of acetylation of proteins in bacterial metabolism, transcription, and other processes.

## MATERIALS AND METHODS

### Chemicals

HPLC-grade acetonitrile and water were obtained from Burdick & Jackson (Muskegon, MI). Reagents for protein chemistry including iodoacetamide, dithiothreitol (DTT), ammonium bicarbonate, formic acid (FA), and urea were purchased from Sigma Aldrich (St. Louis, MO). Sequencing grade trypsin was purchased from Promega (Madison, WI). HLB Oasis SPE cartridges were purchased from Waters (Milford, MA).

### Strains, media, and growth conditions

All strains used in this study are listed in **Table 2 (64-66)**. *E. coli* strains were aerated at 225 rpm in TB7 (10 g/L tryptone buffered at pH 7.0 with 100 mM potassium phosphate) supplemented with 0.4% glucose with a flask-to-medium ratio of 10:1 at 37°C. Derivatives were constructed by moving the appropriate deletions from the Keio collection (67) by generalized transduction with P1kc, as described (68). Kanamycin cassettes were removed, as described (67). Plasmids carrying known and putative GNAT family members were isolated from the ASKA collection (69) and transformed into the indicated strains. Mutagenesis of the plasmids was performed via Quikchange II Site-Directed Mutagenesis Kit (Agilent Technologies) using primers listed in **Table 3**. To maintain plasmids, chloramphenicol was added to a final concentration of 25 μg/mL. To induce GNAT expression from the pCA24n plasmid, IPTG (Isopropyl β-D-1-thiogalactopyranoside) was added to a final concentration of 50 μM or 100 μM.

**Table 2.**
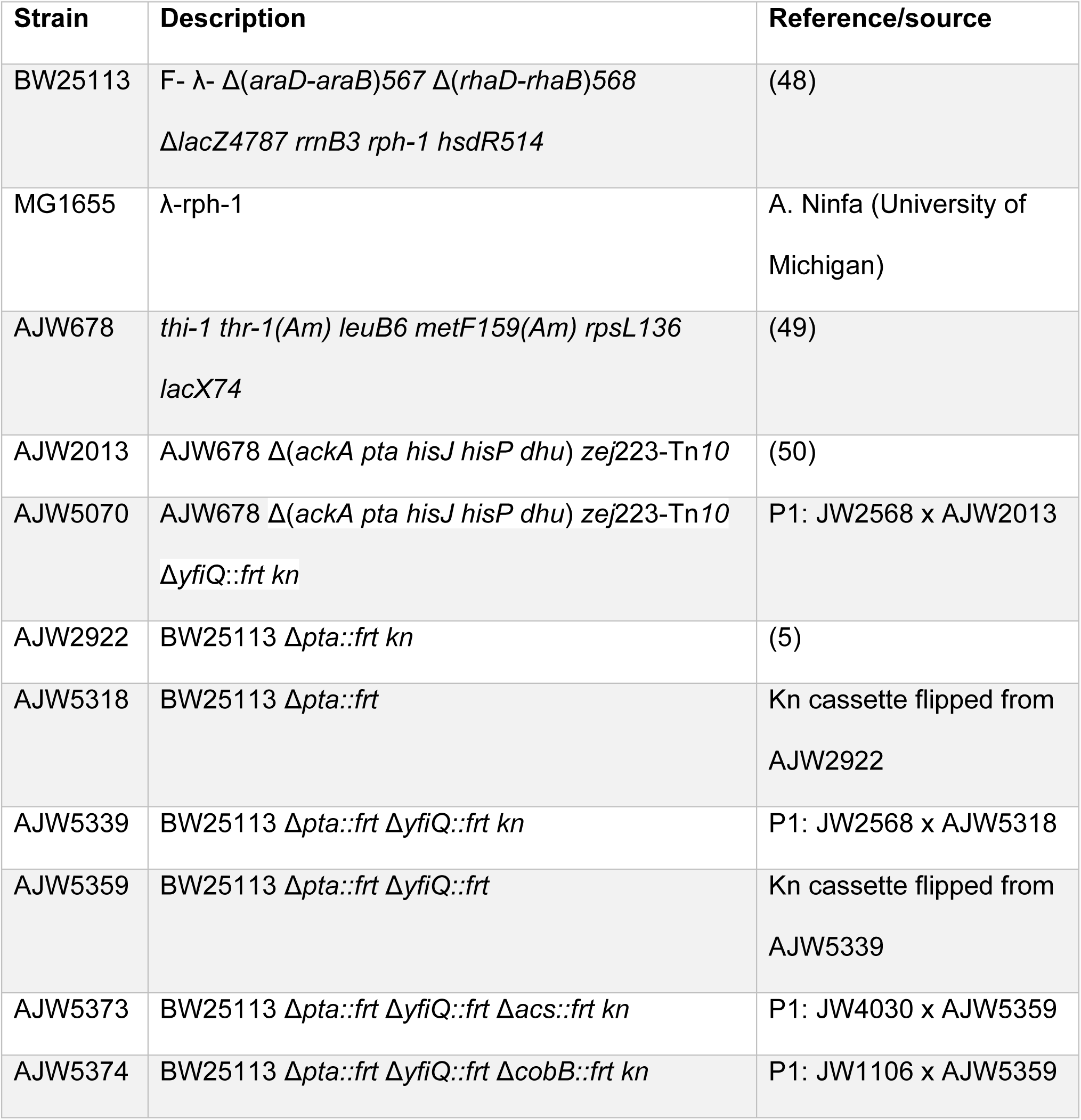
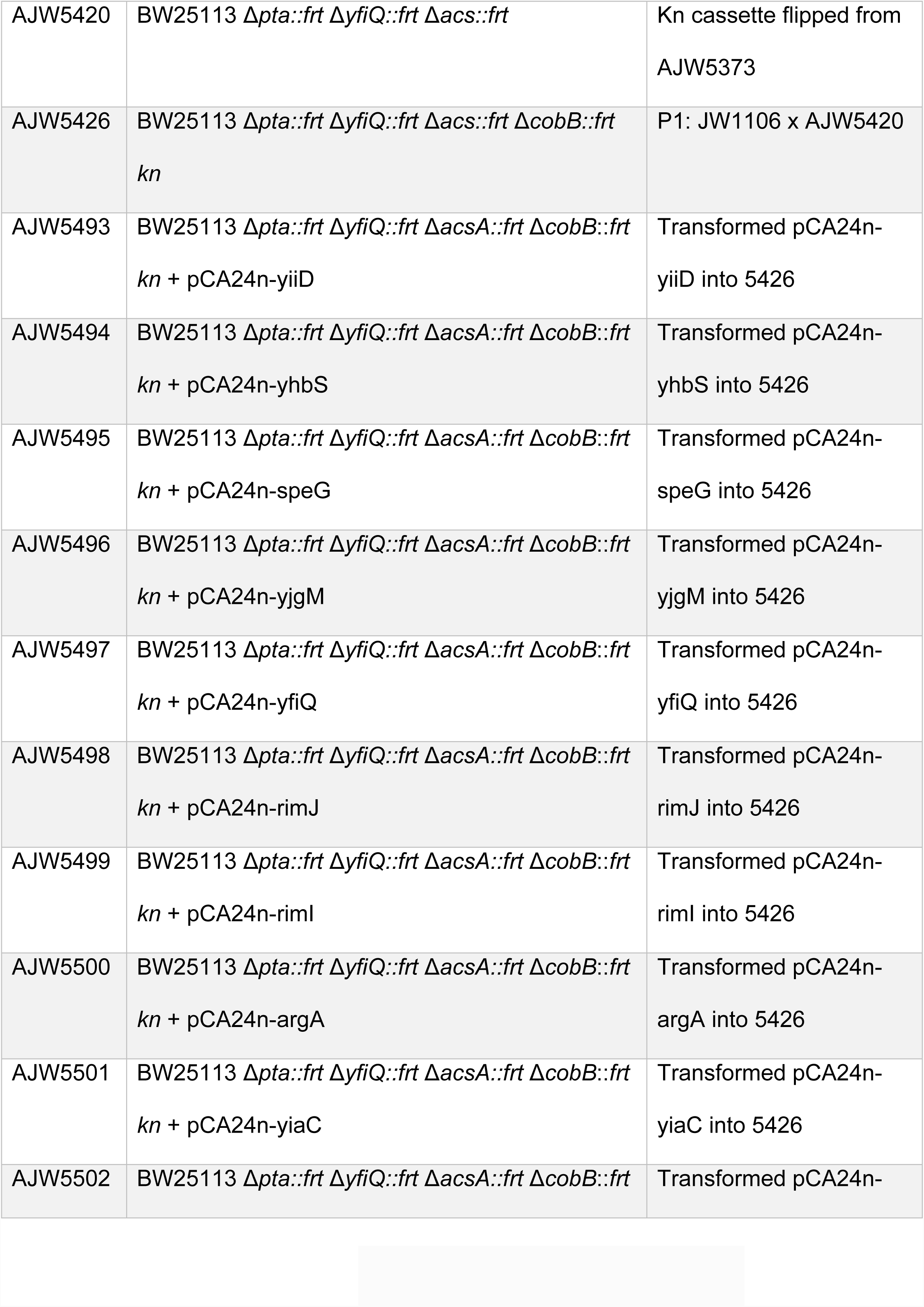
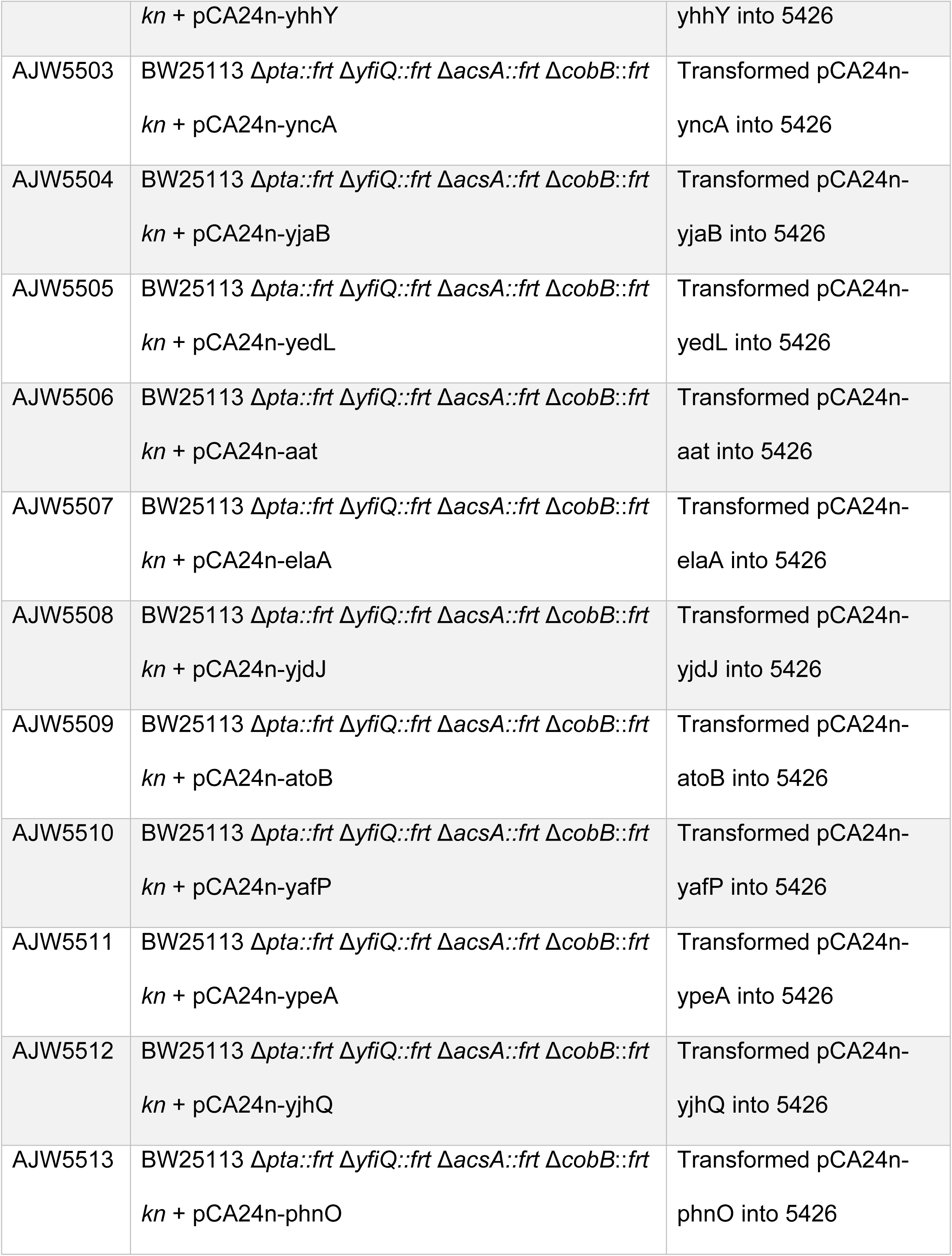
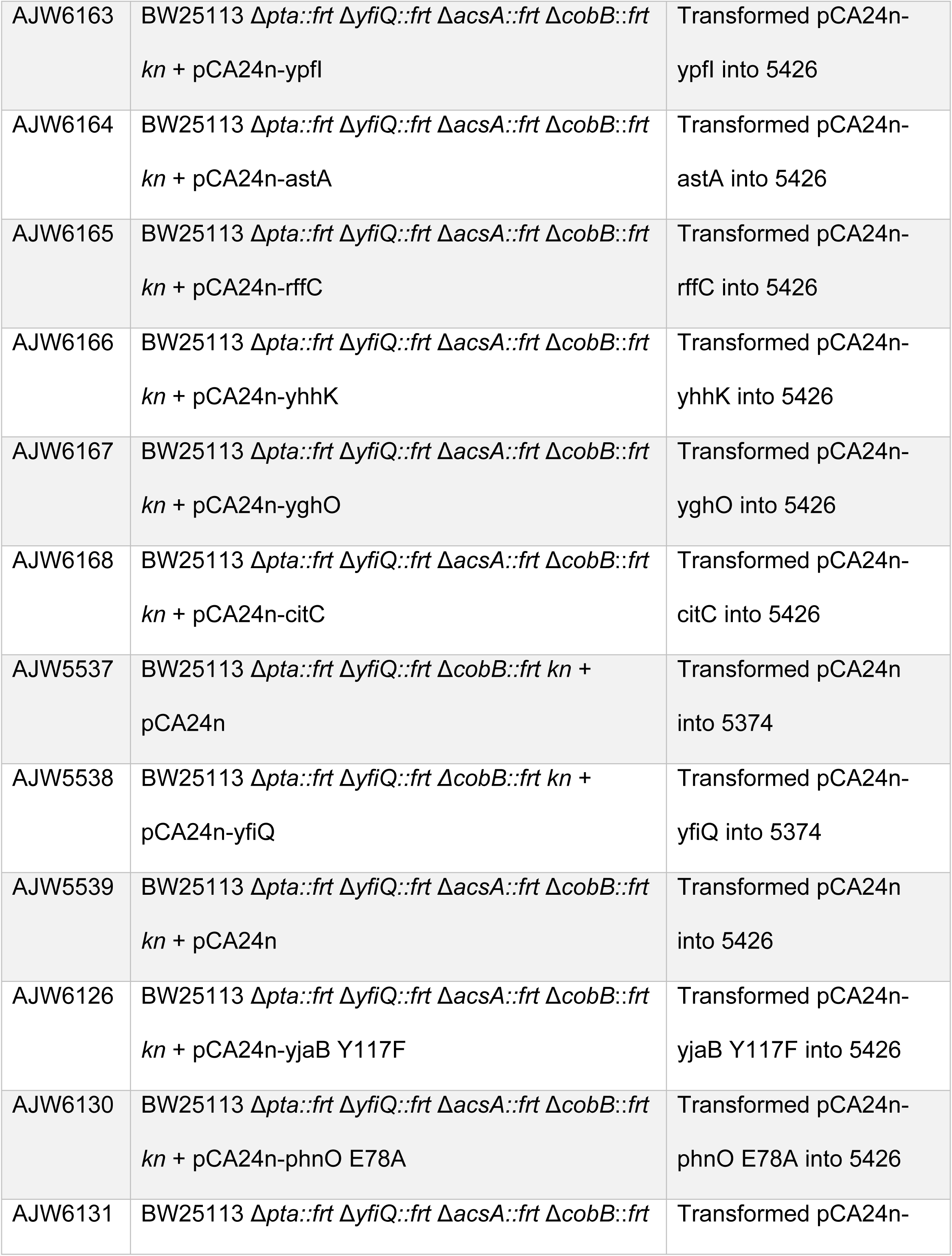
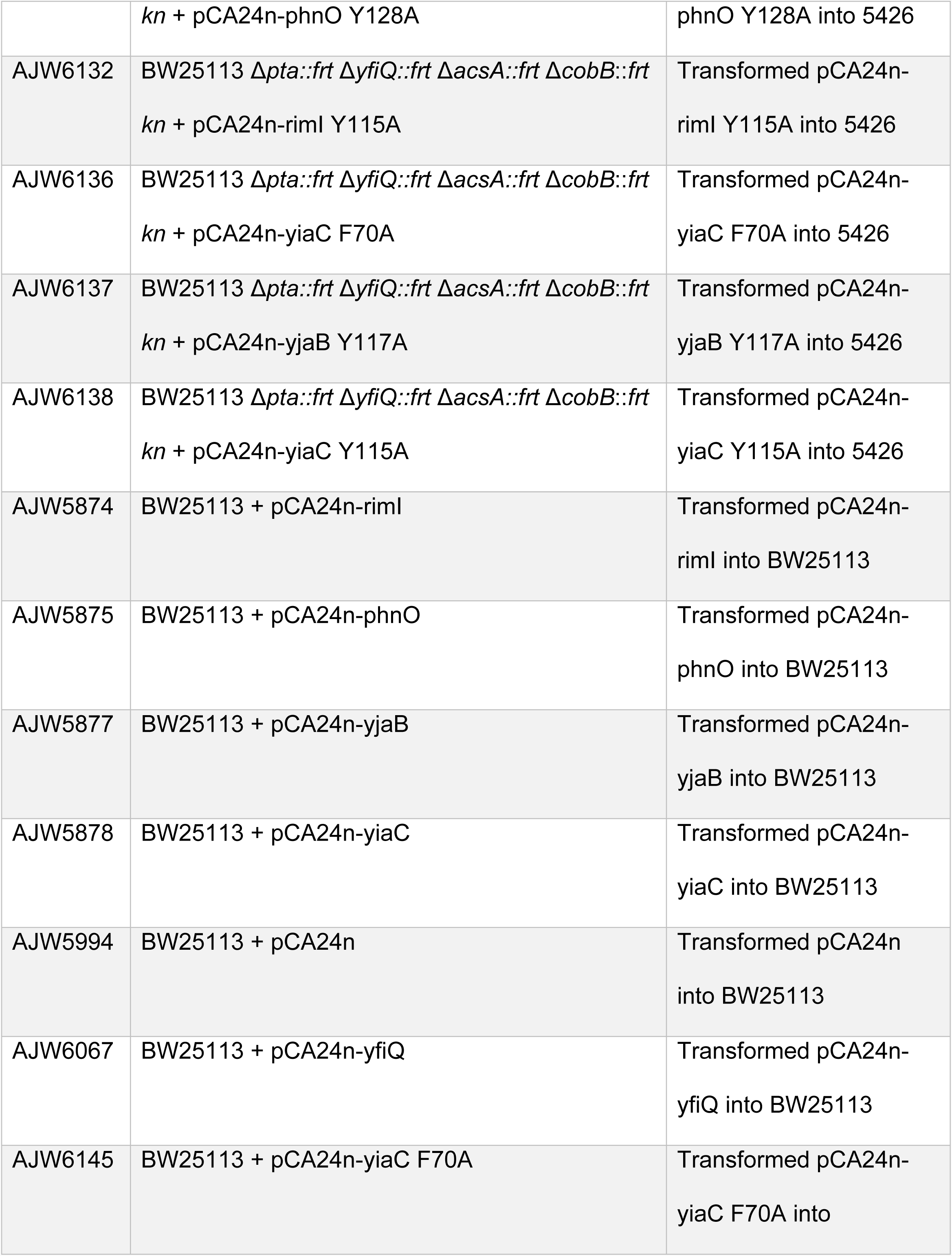
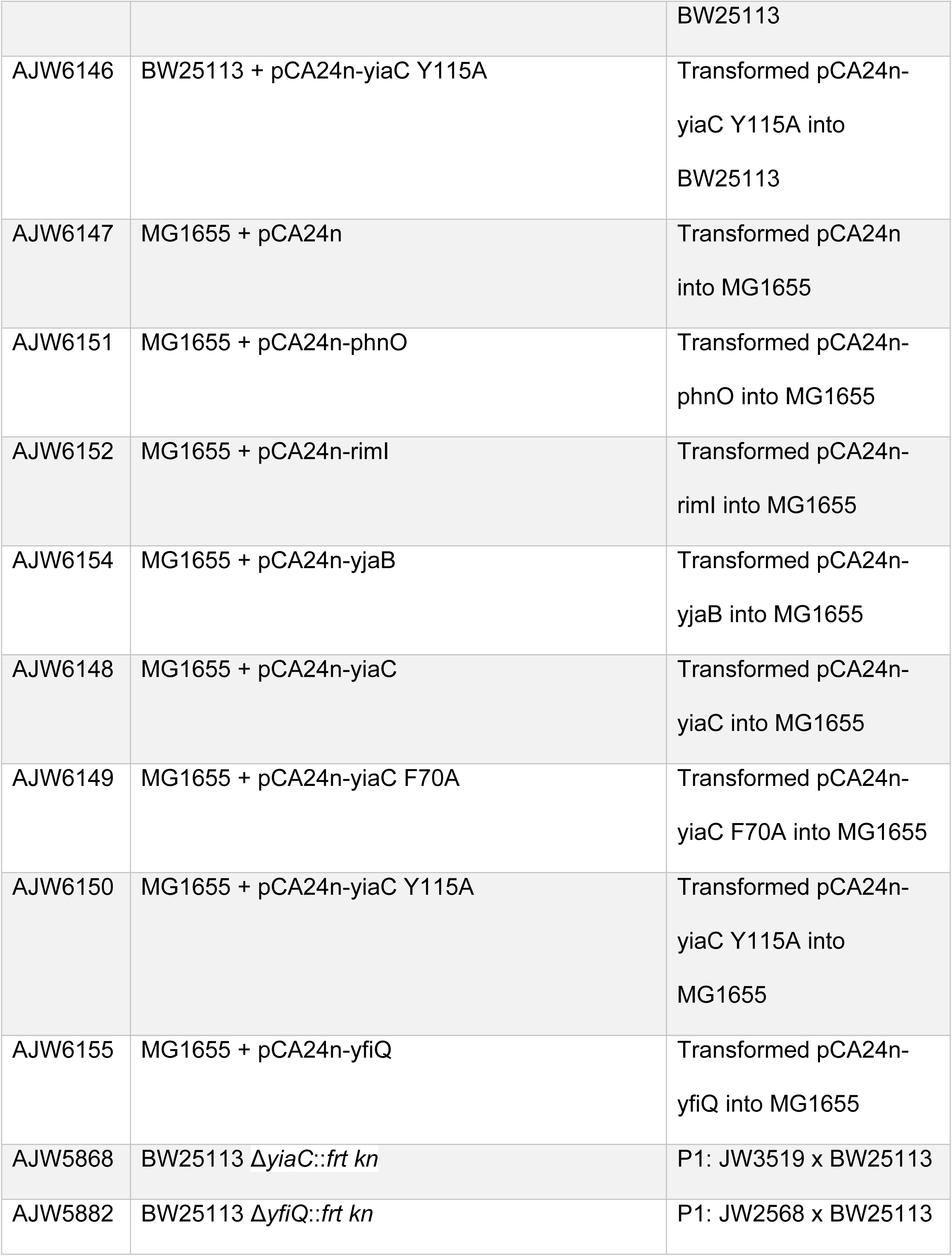
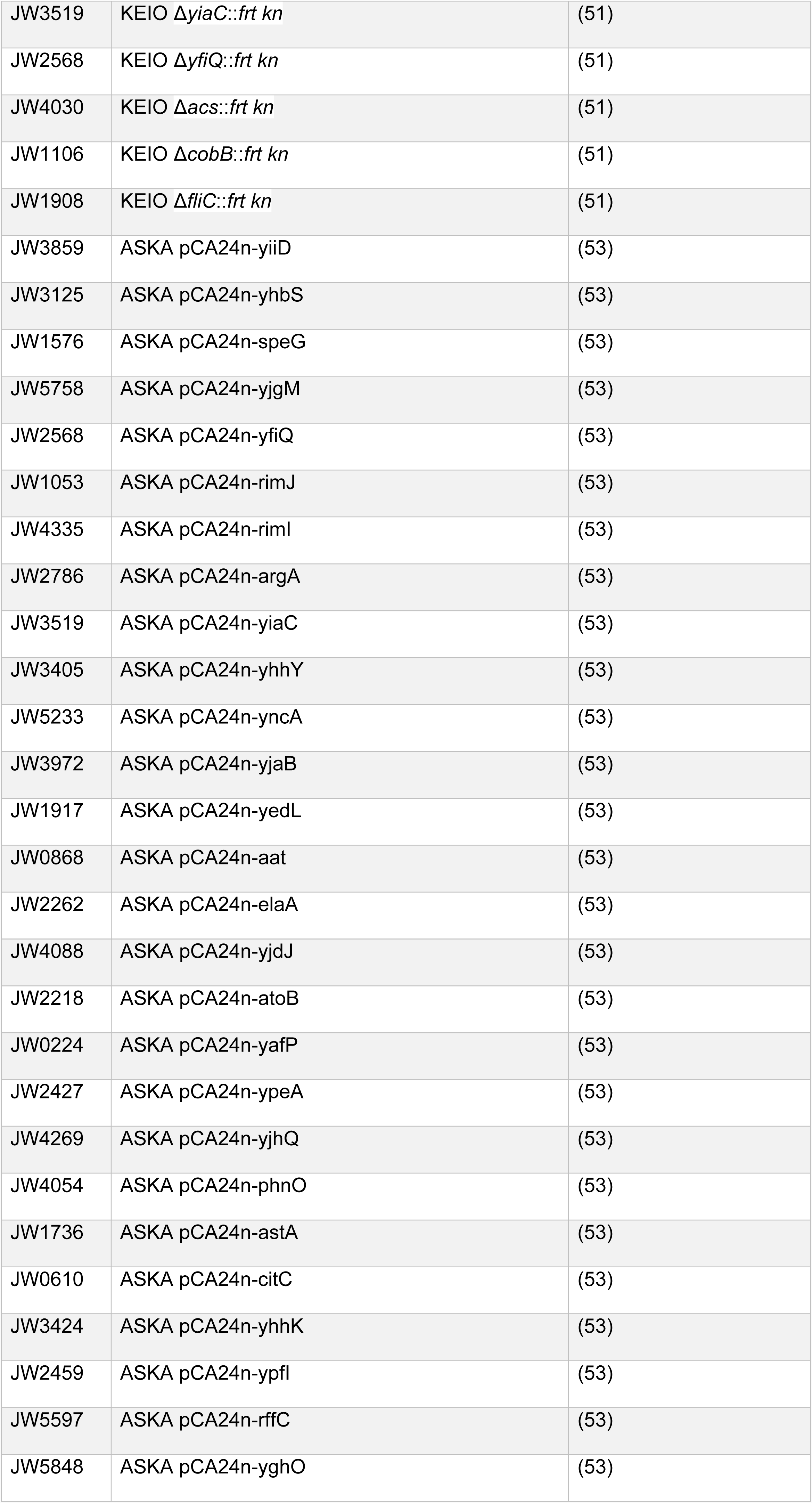
Bacterial Strains and Plasmids

**Table 3.**
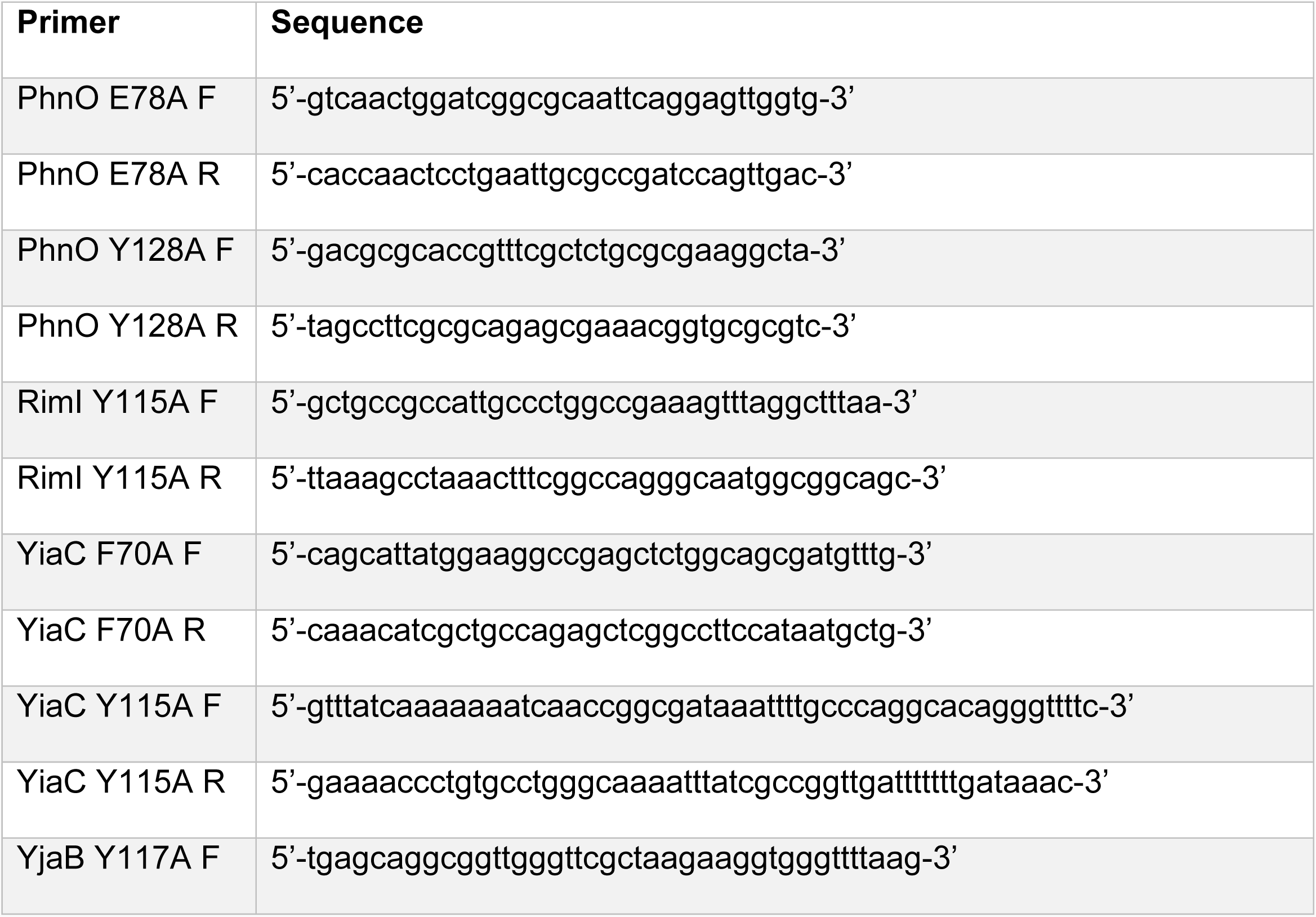
Primers used in this study

### Western blot analysis of protein acetylation and detection of His-tagged proteins

*E. coli* cells were aerated at 37°C in TB7 supplemented with 0.4% glucose for 10 hours unless otherwise noted. When necessary, chloramphenicol was added to a final concentration of 25 μg/mL, while IPTG was added to a final concentration of 50 μM. Bacteria were harvested by centrifugation and lysed using Bugbuster protein extraction reagent (Novagen, Merck Millipore, Billerica, MA). The amount of cell lysate loaded on the gel was normalized to the total protein concentration, as determined by the bicinchoninic acid (BCA) assay (*Thermo Scientific Pierce, Waltham,* MA). Proteins were separated by sodium dodecyl sulfate polyacrylamide gel electrophoresis (SDS-PAGE) and normalization was verified by Coomassie staining. Protein acetylation was determined using a rabbit polyclonal anti-acetyllysine antibody (Cell Signaling, Danvers, MA) at a dilution of 1:1000, as described previously (5, 23).

His-tagged proteins were detected with crude lysates. One milliliter of 1 OD_600_ culture was harvested, pelleted, and resuspended in 200 μl sample loading buffer. The samples were boiled for 10 minutes, and loaded directly onto SDS-PAGE gels. His-tagged proteins were detected with mouse anti-His tag (27E8) antibody at a dilution of 1:1000 (Cell Signaling, Danvers, MA) and an anti-mouse IgG HRP linked antibody at a dilution of 1:2000 (Cell Signaling, Danvers, MA).

### Cell lysis, proteolytic digestion of protein lysates, and affinity enrichment of acetylated peptides

For mass spectrometric analysis, bacteria were cultivated as described for western blot analysis. We then processed isolated frozen bacterial pellets from the gutted strains carrying vector control (AJW5426) or one of the 5 KAT candidates *(i)* RimI (AJW5499), *(ii)* YiaC (AJW5501), *(iii)* YjaB (AJW5504), *(iv)* PhnO (AJW5513), as well as the known KAT YfiQ (AJW5497). Each of the strains was processed as 3 biological replicates. Cell pellets of the indicated strains were suspended in 6 mL of PBS and centrifuged at 4°C, 15,000 g for 20 min. The firm cell pellet was suspended and denatured in a final solution of 6 M urea, 100 mM Tris, 75 mM NaCl, and the deacetylase inhibitors tricostatin A (1 mM) and nicotinamide (3 mM). Samples were sonicated on ice (5× each for 15 sec), cellular debris was removed by centrifugation, and the supernatants were processed for proteolytic digestion. Lysates containing 1.5 mg of protein were reduced with 20 mM DTT (37°C for 1 h), and subsequently alkylated with 40 mM iodoacetamide (30 min at RT in the dark). Samples were diluted 10-fold with 100 mM Tris (pH 8.0) and incubated overnight at 37°C with sequencing grade trypsin (Promega) added at a 1:50 enzyme:substrate ratio (wt/wt). In parallel, separate 1.5 mg protein aliquots were digested using endoproteinase Glu-C (Roche, Indianapolis, IN) by adding Glu-C at a 1:50 protease to substrate protein ratio (wt:wt), and incubating overnight at 37 °C. Subsequently, samples were acidified with formic acid and desalted using HLB Oasis SPE cartridges (Waters) (Keshishian *et al*., 2007). Proteolytic peptides were eluted, concentrated to near dryness by vacuum centrifugation, and suspended in NET buffer (50 mM Tris-HCl, pH 8.0, 100 mM NaCl, 1 mM EDTA). A small aliquot of each protein digestion (∼ 10 μg) was saved for protein-level identification and quantification. The remaining proteolytic peptide samples were used for affinity purification of acetylated peptides (K^ac^).

Acetylated peptides were enriched using 1/4 tube of anti-acetyl lysine antibody-bead conjugated ‘PTMScan Acetyl-Lysine Motif [Ac-K]’ Kit (Cell Signaling Technologies) for each of the 1 mg protein lysate samples according to the manufacturer’s instructions. Prior to mass spectrometric analysis, the acetylated peptide enrichment samples were concentrated and desalted using C-18 zip-tips (Millipore, Billerica, MA).

### Mass Spectrometric Analysis

Samples were analyzed by reverse-phase HPLC-ESI-MS/MS using the Eksigent Ultra Plus nano-LC 2D HPLC system (Dublin, CA) combined with a cHiPLC System, which was directly connected to a quadrupole time-of-flight SCIEX TripleTOF 6600 mass spectrometer (SCIEX, Redwood City, CA) (23). After injection, peptide mixtures were transferred onto a C18 pre-column chip (200 µm x 6 mm ChromXP C18-CL chip, 3 µm, 300 Å, SCIEX) and washed at 2 µl/min for 10 min with the loading solvent (H_2_O/0.1% formic acid) for desalting. Subsequently, peptides were transferred to the 75 µm x 15 cm ChromXP C18-CL chip, 3 µm, 300 Å, (SCIEX), and eluted at a flow rate of 300 nL/min with a 3 h gradient using aqueous and acetonitrile solvent buffers (23).

*Data-dependent acquisitions:* To build a spectral library for protein-level quantification, the mass spectrometer was operated in data-dependent acquisition (DDA) mode where the 30 most abundant precursor ions from the survey MS1 scan (250 msec) were isolated at 1 m/z resolution for collision induced dissociation tandem mass spectrometry (CID-MS/MS, 100 msec per MS/MS, ‘high sensitivity’ product ion scan mode) using the Analyst 1.7 (build 96) software with a total cycle time of 3.3 sec as previously described (5, 23, 70).

*Data-independent acquisitions:* For quantification, all peptide samples were analyzed by data-independent acquisition (DIA, e.g. SWATH) (71), using 64 variable-width isolation windows (5, 72, 73). The SWATH cycle time of 3.2 sec included a 250 msec precursor ion scan followed by 45 msec accumulation time for each of the 64 variable SWATH segments.

### Mass Spectrometric Data Processing and Bioinformatics

Data-independent acquisitions (DIA) from acetyl-peptide enrichments were analyzed using the PTM Identification and Quantification from Exclusively DIA (PIQED) workflow and software tool (24). PIQED uses multiple open source tools to accomplish automated PTM analysis, including Trans-Proteomic Pipeline (74), MS-GF+ (75), DIA-Umpire (76), mapDIA (77), and Skyline (78). Relative quantification of acetylation levels from putative KATs versus VC in biological triplicates was used to determine fold-changes (see Supplemental **Table S1A**). Sites were called regulated when FDR<0.01 and fold change >4. **Supplemental Tables S4A** and **S4B** contain details of all acetylated peptide identifications from the trypsin digestion and GluC digestion experiments, respectively. **Supplemental Tables S4C** and **S4D** contain the unfiltered quantification results of all acetylation sites quantified from trypsin digestion and GluC digestion experiments, respectively. **Supplemental Table S4E** contains details of all proteins identified from ProteinPilot used for spectral library building and protein-level quantification. **Supplemental Table S1D** gives the protein-level changes due to YfiQ overexpression. **Supplemental Table S2B** shows the site-level acetylation changes in proteins related to glycolysis shown in **Fig. 5**.

### Data Availability

All raw mass spectrometry data files are available from public repositories (MassIVE ID number: MSV000082411 and password: kitkats, and ProteomeXchange: PXD009940). The MassIVE repository also includes the supplemental tables, and details of proteins and peptides that were identified and quantified by mass spectrometric analysis. Skyline files containing spectral libraries and chromatograms of raw data quantification are available on Panorama (https://panoramaweb.org/KAT.url, email login: and password: ^x3GfCJh).

### KAT sequence alignment and homology modelling

A multiple sequence alignment containing each *E. coli* KAT sequence (YfiQ UniProt ID: P76594, YiaC UniProt ID: P37664, YjaB UniProt ID: P09163, RimI UniProt ID: P0A944, and PhnO UniProt ID: P16691) was generated using the multiple alignment Clustal W function and manually modified in BioEdit (79). Only the GNAT domain (residues 726-881) of YfiQ was used in the sequence alignment. The final alignment figure was prepared using ESPript 3.0 (http://espript.ibcp.fr) (80). Homology models for YiaC, PhnO, and the GNAT domain of YfiQ were constructed using the ModWeb server (https://modbase.compbio.ucsf.edu/modweb/) (81) with the slow restraint selected for model generation. The models with the highest ModPipe Quality Score (MPQS), the lowest Discrete Optimized Protein Energy (zDOPE) value, a GA341 model score that was closest to one, and the highest sequence identity were chosen for further analysis. The templates used for each of the final homology models were PDB ID: 2kcw for YiaC, PDB ID: 1z4e for PhnO and PDB ID: 4nxy for the GNAT domain of YfiQ.

### Motility-related assays

Cultures were grown in TB (10 g/L tryptone, 5 g/L NaCl) at 37°C to exponential phase (0.3 – 0.5 OD_600_) were normalized to 0.3 OD_600_ and a 5 μl aliquot was spotted onto the surface of a tryptone agar plate (10 g/L tryptone, 5 g/L NaCl, 2 g/L agar). The diameter of the spot was measured hourly. For strains harboring plasmids, IPTG and chloramphenicol were added to both the growth medium and agar plates at a final concentration of 50 μM and 25 μg/mL, respectively.

## Supplemental Figure Legends

**FIGURE S1. A strain lacking known mechanisms of acetylation contains residual lysine acetylation.** Wild-type (WT) *E. coli* (strain AJW678) and an isogenic Δ*ackA pta yfiQ* mutant were aerated in TB7 supplemented with 0.4% glucose for 8.5 hours. Whole cell lysates were analyzed (left panel) by Coomassie blue-stained SDS-polyacrylamide gel electrophoresis to ensure equivalent loading and (right panel) by anti-acetyllysine western blot.

**FIGURE S2. Overexpressing YfiQ results in increased acetylation.** BW25113 Δ*pta yfiQ cobB* cells (Acs+) or Δ*pta yfiQ acs cobB* (Acs-) were transformed with pCA24n-YfiQ or the empty vector (VC). The resultant strains were aerated in TB7 supplemented with 0.4% glucose and 25 μg/ml chloramphenicol for 10 hours. IPTG was added to a final concentration of 50 μM where indicated. Whole cell lysates were analyzed (bottom panel) by Coomassie blue-stained SDS-polyacrylamide gel to ensure equivalent loading and (top panel) by anti-acetyllysine western blot. The acetylated Acs band is indicated by an asterisk (*). Note, leaky expression of YfiQ results in acetylation of Acs in the absence of IPTG.

**FIGURE S3. Five GNAT family members reproducibly alter lysine acetylation patterns.** The gutted strain (BW25113 Δ*pta yfiQ acs cobB*) was transformed with the pCA24n vector control (VC) or pCA24n containing the indicated genes under control of an IPTG inducible promoter (53). The resultant strains were aerated in TB7 supplemented with 0.4% glucose, 50 μM IPTG, and 25 μg/ml chloramphenicol for 10 hours. Whole cell lysates were analyzed (left panels) by Coomassie blue-stained SDS-polyacrylamide gel to ensure equivalent loading and (right panels) by anti-acetyllysine western blot. Triplicate biological samples of each strain are shown (designated A, B, and C when not loaded next to one another).

**FIGURE S4. The hydroxyl group of tyrosine 117 is required for the KAT activity of YjaB.** The gutted strain (BW25113 Δ*pta yfiQ acs cobB*) was transformed with the pCA24n vector control, pCA24n carrying the wild-type YjaB or the indicated YjaB mutants. The resultant strains were grown in TB7 supplemented with 0.4% glucose, 100 μM IPTG, and 25 μg/mL chloramphenicol for 8 hours. Crude lysates harvested after 4 hours were analyzed for expression of the KAT proteins. Whole cell lysates harvested after 8 hours were analyzed for acetylation. Coomassie stained SDS-PAGE gels (left) served as loading controls for anti-His (top right) and an anti-acetyllysine (bottom right) western blots.

**FIGURE S5. Specificity of novel KATs**. (A) Acetyl-lysine sites that were statistically increased due to KAT overexpression were compared with previously reported *ackA*-regulated acetylation sites. (B and C) Motif analysis of the sites regulated by YfiQ and YiaC shows no preference for primary amino acid sequence surrounding the modification site.

**FIGURE S6. YiaC requires catalytic activity to inhibit migration in MG1655.** Cultures were grown overnight in TB medium supplemented with chloramphenicol and 50 μM IPTG. 5 μL of each normalized culture was spotted on low percentage TB plates supplemented with chloramphenicol and 50 μM IPTG. The diameter of the cell spot was measured hourly. (A-C) Growth curves of wild-type *E. coli* strain MG1655 strains carrying the indicated plasmids. (D-F) Hourly migration of wild-type *E. coli* strain MG1655 strains carrying pCA24n encoding the indicated plasmids. (G) Representative image of a motility plates with MG1655 carrying the indicated plasmids.

**Supplemental Table Legends**

**Table S1. Quantification of acetylation and protein level changes upon overexpression of KATs via mass spectrometry.**

**Table S2. Translation-related and metabolic proteins with significant increases in acetylation.**

**Table S3. Conservation of RimI, PhnO, YiaC, and YjaB across bacterial phylogeny.**

**Table S4. All acetylated peptides detected and unfiltered quantification results of all acetylation sites from trypsin and GluC digestions.**

## Acknowledgments

We sincerely thank Drs. George Gassner, Wayne Anderson and Ekaterina Filippova for their helpful discussions.

This project was funded by R01 GM066130 (NIGMS to AJW), R01 AI108255 (NIAID subcontract to AJW), R01 AI108255 (NIAID to BS), a Center for Computing for Life Sciences (CCLS) grant and Startup funds (from SFSU to MLK), and JGM was supported by a (T32 AG000266, NIH to Campisi).

